# Track to Combat Wheat Stem Rust (*Puccinia graminis* f. sp. *tritici*) Races: Pathogenicity Spectrum, Tempo-Spatial Dynamics, and Impacts on Irrigated Wheat in Ethiopia under Climate Change

**DOI:** 10.1101/2025.09.14.676115

**Authors:** Nurhussein Seid, Kitessa Gutu, David P. Hodson, Yoseph Alemayehu, Zemedkun Alemu, Sileshi Getahun, Jemal Tolla, Ayele Badebo, Mohammed Yesuf

## Abstract

1.

Wheat stem rust (Puccinia graminis f. sp. tritici) is a major global threat to wheat production, driven by rapid shifts in virulence and race diversity. This study aimed to identify physiological races of stem rust pathogens in Ethiopia’s irrigated wheat-growing areas. Surveys and race analyses were conducted during the 2020/21–2023/24 large-scale wheat expansion periods, alongside a systemic review of rust dynamics from 2012 to 2022. Results revealed significant shifts in stem rust races, with an increasing dominance of virulent races over the last decade. TKTTF dominated from 2012 to 2016, succeeded by TTTTF in 2017, and TKKTF in 2019–2020. By 2021–2022, TTKTT and TTKTF races were prevalent, affecting 90% of wheat fields. From 2020 to 2024, five major races were identified: TTKTT, TTKTF, TTTTF, TKKTF, and TKTTF, with TTKTT, a virulent Ug99 mutant, emerging as the most dominant. This race exhibits a 95% virulence spectrum, overcoming resistance genes such as Sr24, widely used in commercial cultivars. The study highlights the urgent need to prioritize resistance breeding against virulent races, particularly in irrigated wheat regions, which serve as hotspots for pathogen evolution. Strategic rust intervention and improved screening are essential for protecting both irrigated and rain-fed wheat crops, ensuring sustainable wheat production in Ethiopia and the globe under climate change.

**Abstract:** Wheat stem rust (Puccinia graminis f. tritici) is a significant concern for farmers and agricultural communities around the world. It is essential to monitor the changes in virulence among these infections, as understanding these shifts can help prevent unexpected epidemics that impact livelihoods and food security.

**Objective:** This study seeks to compassionately uncover the physiological races of stem rust pathogens present in irrigated wheat-growing regions of Ethiopia, recognizing the vulnerability of local farmers to these challenges.

**Materials and Methods:** We conducted thorough surveys and surveillance, accompanied by detailed physiological and genetic race analyses. The insights gained from this research will be instrumental in guiding the large-scale demonstration and expansion of irrigated wheat from 2020/21 to 2023/24. Furthermore, we conducted comprehensive reviews of the dynamics of stem and yellow rust races affecting wheat production in Ethiopia over the past decade, from 2012 to 2022.

**Results and Discussion:** Our findings reveal that changes in stem rust pathogen races, with a wider pathogenicity spectrum, are prevalent in both rain-fed and irrigated wheat-growing areas of Ethiopia. The rise of virulent races in affected fields has understandably raised concerns among farmers, with frequencies reaching up to 50%. Over the last ten years, the dynamics of wheat rust races illustrate a troubling shift toward more virulent strains. For example, the TKTTF race, virulent to resistance gene 85, was predominant from 2012 to 2016 but was subsequently replaced in 2017 by the more aggressive TTTTF race, which is harmful to gene 90. In 2018, we observed that TTTTF, along with TKTTF and TKKTF (virulent to gene 80), became equally common, creating additional challenges for our farming communities. The TKKTF race held dominance in 2019 and 2020, while from 2021 to 2022, TTKTT (virulent to gene 95) and TTKTF (virulent to gene 85) became more widespread, affecting around 90% of wheat fields.During the years 2021/22 to 2023/24, we carefully collected and analyzed 51 field rust samples to understand the situation better. We identified seven Pgt pathotypes in irrigated wheat in the 2021/22 season: TTKTT, TTKTF, TKKTF, TTRTF, TTTTF, TKTTF, and TKPTF. The emergence of the TTKTT race, a pathogenic Ug99 mutant, particularly in regions like Jimma, Buno Bedale, West Arsi, and East Shoa, is particularly concerning for the local farmers who depend on stable crops. In 2020/21, five stem rust races TTKTT, TTKTF, TTTTF, and TKTTF were identified, with TTKTF and TTKTT showing dominance in geographic distribution. Many popular varieties are now susceptible to these races, adding to the anxiety of farmers who rely on these crops for their livelihoods. The TTKTT race has been particularly alarming, with a 49% prevalence in Ethiopia for the first time, and it has shown varying reactions ranging from susceptible to moderately susceptible among different wheat types. Importantly, the resistance gene Sr24, which many farmers have relied upon in their cultivars, has unfortunately been rendered ineffective against this race.

**Conclusion:** To develop durable resistance, it is crucial to introduce a transgene cassette containing five resistance genes into bread wheat as a single locus within the wheat production area. Integrating robust resistance genes from wild grass relatives with modern scientific breakthroughs is essential. Consequently, breeding programs should focus on identifying additional sources of resistance to counter the more virulent races of the pathogen. Special attention must be given to irrigated wheat production, which harbors virulent and genetically diverse races, threatening the belg and main season wheat crops, as well as the green bridge. The background data collected from this study will aid in strategic rust intervention, screening, and guidance for resistance breeding among wheat breeders and seed technology multiplication units in the area addressing the climate change.

## 2. Introduction

Ethiopia is one of the leading producers of wheat in sub-Saharan Africa. With an annual production of 4.5 million metric tons and cultivation over nearly 1.7 million hectares, wheat accounts for 13.49% of Ethiopia’s cropland. However, the country’s average wheat yield, 2.8 t/ha (CSA, 2019), is lower than the global average of 3.43 t/ha (FAO, 2018). According to Abebe *et al*. (2012), wheat production losses in Ethiopia are primarily caused by biotic, abiotic, and socioeconomic constraints. Among these, diseases, insect pests, and weeds represent the most significant biotic factors limiting wheat production (Abebe *et al*., 2012).

The most critical diseases reducing wheat productivity in Ethiopia are stem rust (*Puccinia graminis* f. sp. *tritici* Eriks. & E. Henn.), leaf rust (*P. triticina* Eriks.), and stripe rust (*P*. *striiformis* Westend f. sp. *tritici*). Rust pathogens have been the most significant biotic stressors in Ethiopia over the past 50 years, as reported by Getaneh (1996) and Temam (1984). Wheat rust outbreaks have been frequently recorded in the Ethiopian Highlands (Saari and Prescott, 1985). Disease epidemics have been common in both durum and bread wheat varieties, with mean infection incidence rates of 44% for stripe rust (Yr), 34% for stem rust (Sr), and 18% for leaf rust (Lr) (Meyer *et al*., 2021). Stem rust, in particular, is a major biotic challenge to wheat production in Ethiopia (Bekele, 1986).

Stem rust, also known as black rust, is considered the most severe of the wheat rust diseases in Ethiopia. It can lead to complete yield losses over extensive areas during epidemic years (Admassu *et al*., 2012; Denbel *et al*., 2013; Olivera *et al*., 2015). The fungus *Puccinia graminis* f. sp. *tritici* (*Pgt*) causes stem rust. The Ethiopian highlands’ southeastern and central-western regions, where most of the country’s wheat is grown, provide conditions conducive to wheat rust infections almost year-round. Presently, modern commercial wheat cultivars in Ethiopia have demonstrated resistance to leaf rust due to advancements in resistant gene development, such as Lr-17, which provides protection at both the seedling and adult plant growth stages.

Key factors contributing to wheat rust prevalence include favorable weather conditions, continuous wheat production, and irrigation practices. Bimodal rainfall patterns in major wheat-growing regions, such as Bale and Arsi, enable two to three wheat harvests within three seasons annually. This creates a “green bridge,” allowing pathogens to spread between wheat-growing seasons (Zadoks and Bouwman, 1985). Specific areas such as Arsi Negele, Debre Zeit, and Herero have been identified as hotspots for rust outbreaks, with recurring rust issues reported in the Shoa, Arsi, and Bale districts. For example, two severe stem rust epidemics occurred on the variety Enkoy in Arsi and Bale during 1993–1994 (Shank, 1994). Cultivars such as Mamba, Romany B.C., Gara, KKBB, Dashen, and Enkoy also lost resistance to one or more rust diseases during this period (Getaneh, 1996).

Several major epidemics have occurred in recent years, including a significant outbreak of wheat stripe rust in 2010 (Olivera, 2015) and another of stem rust on the Digalu variety in 2013, which caused yield losses of up to 100% and an average loss of around 50% (Olivera *et al*., 2015). With climate change, additional outbreaks are anticipated. The emergence of the TTRTF race in East Africa (2016–2020) has broadened the known geographic distribution of this race to include regions such as Italy (2016), Georgia (2014), Hungary (2017), and Egypt (2015–2016) (Patpour, 2020; Tsegaab *et al*., 2020; Olivera *et al*., 2019; Samar and Szabo, 2018).

Ethiopia has been recognized as a hotspot for the development and spread of new Pgt races, as highlighted by Singh et al. (2006). Studies indicate that Ethiopia’s wheat stem rust populations are highly variable, with virulence spectrums ranking among the most severe globally (Admassu et al., 2009; Gutu et al., 2022). Resistance genes such as Sr24, used globally in commercial wheat cultivars, have lost efficacy due to evolving race patterns. The TTKTT race, for instance, was identified in Ethiopian wheat varieties such as Hulluka, Senate, Shorima, Ogolcho, Hidase, and Danda’a. This race has also been observed in Kenya, Iran, Eritrea, and Sudan (Patpour *et al*., 2016). Similarly, TKTTP and TKKTP races, with virulence toward the Sr24 gene, were reported in Turkey and Tunisia (Nazari *et al*., 2022).

Regular monitoring and identification of wheat stem rust races are crucial for understanding the disease’s impact and tracking shifts in virulence. Such surveys provide essential data on race distribution, emergence of new races, and virulence trends, aiding the development of wheat cultivars with long-lasting resistance (Tesfaye *et al*., 2016). Integrating advanced scientific techniques, such as Pgt RNA interference (RNAi) and small RNAs (sRNAs), into wheat pathology research is vital for maintaining the stability of Ethiopia’s wheat production systems.

## 3. Materials and Methods

### 3.1. Collection of infected tissue samples

A comprehensive review of published articles and research on the monitoring and early warning of rust diseases allowed for the evaluation of the temporal dynamics of the wheat stem rust race. In the primary wheat-growing regions of Ethiopia, infected stem samples (two samples per field) were collected from wheat fields at intervals of 5 to 10 kilometers. The survey focused on wheat fields along accessible major and feeder highways.

To prepare the samples for drying, the stem leaf sheath for *Puccinia graminis tritici* (*Pgt*) and the leaf for *Puccinia striiformis* f. sp. *tritici* (Pst) of infected wheat plants were cut into small pieces measuring 5 to 10 cm in length using scissors, then placed in paper bags. This method helps prevent sample deterioration before analysis (Woldeab et al., 2017). The label on each paper bag included details such as the zone, district, variety, GPS data (altitude, latitude, and longitude), and the date of collection. The samples were then transported to the Cereal Disease Laboratory (CDL) at the University of Minnesota and the Ambo Agricultural Research Centre (AARC) laboratory for further analysis.

#### 3.1.1. Isolation and multiplication of single-pustule isolates

Five seeds of the universally susceptible wheat variety (McNair) were planted in pots with a 10 cm diameter, filled with a 2:1:1 volume mixture of sterilized soil, sand, and manure. In the greenhouse, seedlings were grown at a temperature of 18 to 25 °C and a relative humidity of 98 to 100%.

At the seven-day mark, urediniospores from each field were suspended in the light mineral oil Soltrol 170 (Chevron Phillips Chemical Company, The Woodlands, Texas, United States) and applied to the McNair seedlings (Roelfs et al., 1992). Seven days post-inoculation, a leaf with a single fleck that produced one pustule was selected from the base of the leaves. The remaining seedlings in the container were then trimmed with scissors. To avoid cross-contamination, only leaves from each location bearing a single pustule were wrapped in cellophane bags and secured at the base with a rubber band (Fetch and Dunsmore, 2004).

After two weeks, the urediniospores from each pustule were combined with Soltrol 170 to create a solution. This mixture was used to inoculate new susceptible McNair seedlings, also seven days old, intending to multiply each pustule in a separate pot. Following inoculation, the seedlings were placed in a greenhouse after being exposed to four hours of light and 18 hours of dark incubation at 18 to 22 degrees Celsius. After 14 days, the spores from each pustule were isolated and placed in gelatin capsules before being injected into standard differential lines.

The McNair variety, which lacks known resistance genes for stem rust, was infected with urediniospores sourced from each field and suspended in Soltrol 170 using an atomized inoculator on seedlings that were 7 days old (Roelfs et al., 1992).

#### 3.1.2. Inoculation of wheat stem rust differential lines

In pots with a 10 cm diameter, seedlings of the 20 wheat differential host lines, known for their stem rust resistance genes, were raised alongside the susceptible McNair variety. These differential lines were provided to the Ambo Agricultural Research Centre from the USA and Minnesota’s Cereal Disease Laboratory (CDL).

Each rust isolate was suspended in Soltrol 170 following the previously described methods. The suspension was then sprayed onto the seven-day-old differential seedlings (as listed in Table 1) before planting. After inoculation, the plants were transferred to greenhouse benches where the temperature and relative humidity were maintained at 18–25 °C and 98–100%, respectively (Stubbs et al., 1986) (see Table 1).

**Table 1.** List of wheat stem rust differential lines used for race identification.

| Differential host | Sr genes | Differential host | Sr-genes |
| --- | --- | --- | --- |
| LcSr24Ag | 24 | Combination VII | 17 |
| W2691SrTt-1 | 36 | ISr5-Ra | 5 |
| ISr7b-Ra | 7b | ISr6-Ra | 6 |
| ISr8a-Ra | 8a | W2691Sr9b | 9b |
| CnSSrTmp | Tmp | Vernsteine | 9e |
| Sr31 (Benno)/6*LMPG | 31 | W2691Sr10 | 10 |
| CnS-T-mono-deriv | 21 | BtSr30Wst | 30 |
| Trident | 38 | CnsSr9g | 9g |
| ISr9a-Ra | 9a | ISr11-Ra | 11 |
| ISr9d-Ra | 9d | McNair701 | McN |

Source: Roelfs and Martens, 1988; Jin et al., 2008.

Determination of Pgt race was conducted 14 days after inoculation, using seedling infection types on the differential lines, which were graded on a scale from 0 to 4 (Stakman et al., 1962). Infection type readings of 3 (medium-sized uredia with or without chlorosis) and 4 (large uredia without chlorosis or necrosis) were classified as susceptible. The remaining findings, namely 0 (immune or fleck), 1 (small uredia with necrosis), and 2 (small to medium uredia with chlorosis or necrosis), were regarded as low infection types or resistance reactions. Additionally, characters such as “-” (uredinia somewhat smaller than normal for the infection type) and “+” (uredinia slightly larger than normal for the infection type) were modified to highlight variations (Stakman et al., 1962). The races identified on differential lines were categorized into five subsets for racial classification, as illustrated in Table 2. Each isolate was assigned a five-letter identifier based on its response to the differential lines (Roelfs and Martens, 1988; Jin et al., 2008).

**Table 2.**
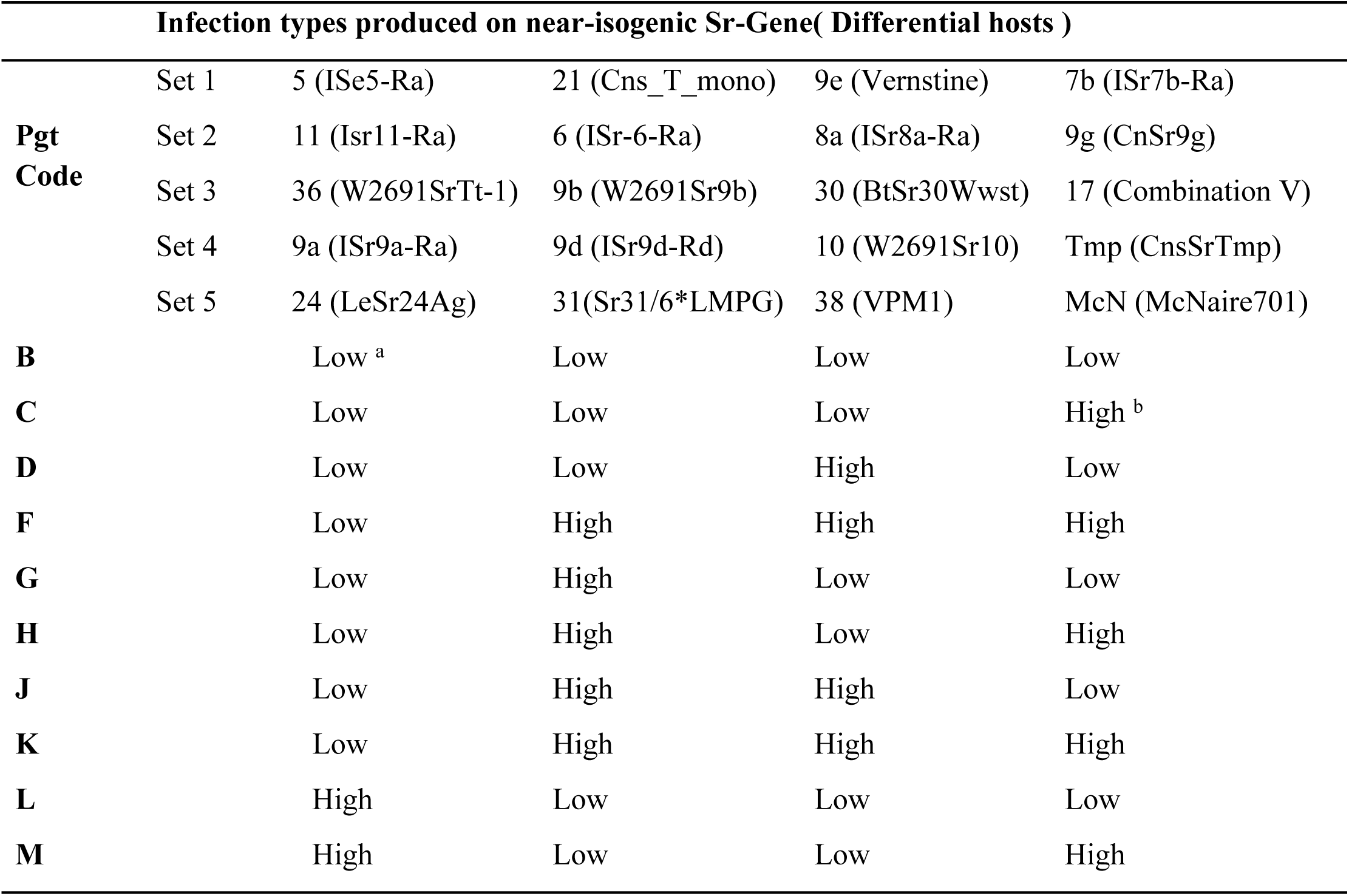
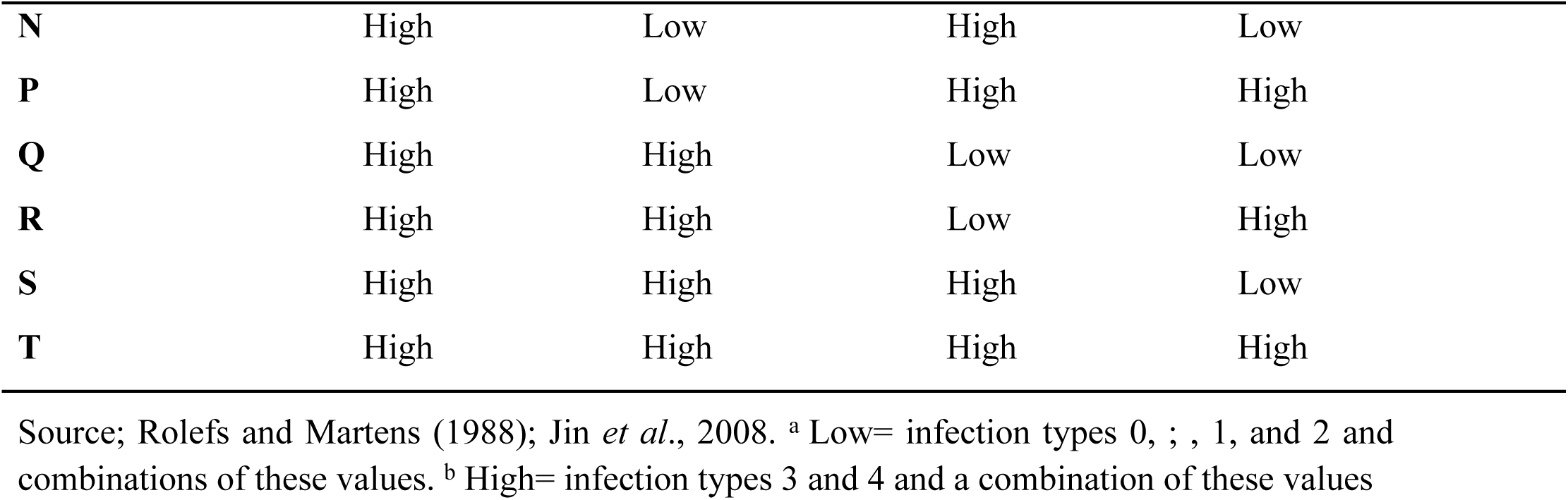
Nomenclature of *Puccinia graminis* f. sp. *tritici* based on 20 differential wheat host lines Infection types produced on near-isogenic.

#### 3.1.3. Inoculation of differential lines and race determination

Five seeds from each of the twenty wheat stem rust differential lines, which carry known stem rust resistance genes (Sr5, Sr6, Sr7b, Sr8a, Sr9a, Sr9b, Sr9d, Sr9e, Sr9g, Sr10, Sr11, Sr17, Sr21, Sr24, Sr30, Sr31, Sr36, Sr38, SrTmp, and SrMcN), along with a susceptible variety (McNair), were cultivated in 10 cm diameter pots. The susceptible variety, McNair (which lacks any Sr gene), was used to assess the survivability of spores injected into different host plants. To maintain each rust isolate obtained from a single pustule, Soltrol-130 was utilized.

A spore suspension was prepared by mixing 3–5 mg of urediniospores with 1 ml of Soltrol-130 and then inoculating it onto the differential seedlings. Fourteen days after inoculation, stem rust infection types (ITs) were evaluated using a scale from 0 to 4, as described by Stakman et al. (1962). Race identification followed a five-letter race code nomenclature system according to Roelfs and Marten (1988), Jin et al. (2008), and Hei et al. (2018). In this system, the differential lines are categorized into five subsets in the following order:

i. Sr5, Sr21, Sr9e, Sr7b
ii. Sr11, Sr6, Sr8a, Sr9g
iii. Sr36, Sr9b, Sr30, Sr17
iv. Sr9a, Sr9d, Sr10, SrTmp
v. Sr24, Sr31, Sr38, SrMcN (Table 1).

An isolate that produces a low infection type on the four lines in a set is assigned the letter ‘B’, while a high infection type is assigned the letter ‘T’. Thus, if an isolate produces a low infection type (resistant reaction) on all 20 differential lines, the race could be designated with a five-letter race code ‘BBBBB’. Conversely, an isolate producing a high infection type would be designated as ‘TTTTT’. Race analysis was conducted using descriptive statistics.

The inoculated seedlings were placed in a dew chamber under complete darkness for 18 hours at a temperature of 18 to 22 °C and relative humidity of 98 to 100%. Fine distilled water droplets were used to hydrate the plants. After removal from the chamber, plants were exposed to four hours of fluorescent light to create an environment conducive to infection, followed by a drying period of about two hours to allow the dew to evaporate. Following inoculation, the plants were moved to benches in a greenhouse, where the environment was controlled to maintain a 12-hour photoperiod, a temperature range of 18 to 25 °C, and a relative humidity (RH) range of 60 to 70% (Stubbs et al., 1986).

After 7–10 days of inoculation, flecks or chlorosis became visible on the leaves. Leaves with a single fleck that developed into a single pustule (single uredinial isolate) were selected from the leaf base, while the remaining seedlings in the pots were removed with scissors. Only leaves with single pustules from each location were separately covered with Cellophane bags and secured at the base with a rubber band to prevent cross-contamination (Fetch and Dunsmore, 2004).

When the pustules appeared after two weeks, the spores from each pustule were isolated and preserved separately in gelatin capsules using a power-operated vacuum aspirator. Following the previously described techniques, a suspension made by combining urediniospores with Soltrol 170 was inoculated on seven-day-old seedlings of the susceptible variety McNair for multiplication of each of the single pustules in separate pots. A single pustule isolate was derived from the urediniospores that originated from a single pustule. For the final race analysis, one isolate was created from each field of wheat (Hei et al., 2018).

### 3.2. Genetic Race Analysis

As a follow-up to the physiological race analysis, dead stem rust samples were collected for genetic race analysis. This was conducted on sample sets gathered from irrigated wheat areas between 2019 and 2022. In these regions, a total of 9, 33, 18, and 34 samples were collected from widely scattered and representative locations. Sharp scissors were used to cut above and below a single pustule to obtain the isolate. The stem inside the sheath tissue was then removed from the side. After collection, the samples were stored in 80% alcohol for seven days. They were subsequently left to dry in the open air for two days. Finally, the samples were delivered in sealed tubes to the Cereals Disease Lab (CDL) in Minnesota for DNA analysis.

### 3.3. Data analysis

The race analysis results from the survey data were examined using descriptive statistics (means) annually across the nation. The dynamics of racial shifts were presented using graphical displays, following the methodology outlined by Gomez and Gomez (1984).

## 4. Results and Discussion

### 4.1. Virulence and physiological races of Stem rust pathogen

During our study, we collected a total of 33 stem rust samples from irrigated areas and sent them to the Ambo Agricultural Research Center’s stem rust laboratory to analyze the distribution and frequency of Puccinia graminis tritici (Pgt) races. Unfortunately, only four of these samples, representing different races, were viable, resulting in a low viability rate of 12.12%. The causes for this low viability remain unidentified but may be attributed to delays in collecting the samples or issues related to the handling of the tested samples.

In the 2019-2020 crop **seasons**, we collected a total of 46 stem rust samples for physiological race analysis at the AARC; 13 samples (28.26%) were not viable, leaving us with 33 viable samples. From this crop season, 28 samples were collected from the Oromia region (specifically from the Sire, Jeju, and Fentale districts), while 19 samples were obtained from the Afar regional state (from the Afambo, Dubti, and Amibara districts).

#### 4.1.1. Overview of Sample Viability and Collection

For the 2020-2021 production year, we sent a total of 51 stem rust samples to the AARC for physiological race analysis to assess the distribution and frequency of Pgt races in irrigated areas. In the following year, 2021-2022, we sent 99 samples for the same purpose. During the 2022-2023 and 2023-2024 production years, we collected and sent 51 live samples from irrigated wheat production areas.

However, we observed that only 21.6% of the samples were viable due to unidentified factors, which is considered quite low for a research year. Additionally, in surveys conducted during 2022, a total of 46 stem rust samples were sent to the AARC for further physiological race analysis. Of these, 28 samples were from the Oromia region (Sire, Jeju, and Fentale districts) and 19 from the Afar regional state (Afambo, Dubti, and Amibara districts) (see Table 3).

**Table 3.** Wheat Stem rust race composition in the irrigated wheat production areas of Ethiopia.

| Year | Live sample | Viability | TTKTF | TTKTT | TKKTF | TTRTF | TTTTF | TKTTF | TKPTF |
| --- | --- | --- | --- | --- | --- | --- | --- | --- | --- |
| 2017/18 | 0 | 0 | 0 | 0 | 0 | 0 | 0 | 0 | 0 |
| 2018/19 | 33 | 12.12 | 25 | 0 | 25 | 0 | 25 | 25 | 0 |
| 2019/20 | 46 | 71.74 | 12.5 | 0 | 24.5 | 0 | 45.5 | 15.2 | 3 |
| 2020/21 | 51 | 21.6 | 36.4 | 4.5 | 0 | 18.2 | 9.1 | 27.3 | 4.5 |
| 2021/22 | 99 | 66.6 | 38.8 | 49 | 6.8 | 0 | 9.1 | 4.5 | 0 |
| 2022/23 | 51 | 43.4 | 66.7 | 33.33 | 0 | 0 | 0 | 0 | 0 |
| 2023/24 | 51 | 42.3 | 66.7 | 22.2 | 11.1 | 0 | 0 | 0 | 0 |

#### 4.1.2. Dominant Physiological Races Identified

The study identified seven physiological races of Pgt: TTKTF, TTKTT, TKKTF, TTRTF, TTTTF, TKTTF, and TKPTF. Among these, TKKTF was the most prevalent, constituting 48.2% of the total samples, followed by TKTTF (33.7%). Less frequent races included TTTTF (4.8%), TTKTT (2.4%), and TTRTF (1.2%). These races showed significant variability in virulence, with many overcoming resistance genes commonly present in commercial wheat cultivars worldwide. For example: TTKTT exhibited a virulence spectrum of 95%, rendering genes such as Sr24 ineffective. TKKTF and TKTTF demonstrated 80% and 85% virulence frequencies, respectively, defeating most known resistance genes, including Sr24 and Sr31.

The highly aggressive TTTTF race eliminated 90% of resistance genes.

Puccinia graminis f.sp. tritici (pgt) comprises seven physiological races, including TTKTT and TTKTF. We identified the following races: TKKTF, TTRTF, TTTTF, TKTTF, and TKPTF. The most prevalent race, TKKTF, was found in 40 samples (48.2%), followed by TKTTF (the Digelu race), which was present in 28 samples (33.7%). The less common races, TTTTT, TTKTT, and TTTTF, were identified in three samples, with occurrences of one (1.2%), two (2.4%), and four (4.8%), respectively. Notably, 80% to 100% of the resistance genes in various host lines were eliminated by these races. Among the detected races, the resistance genes Sr24 and Sr31 proved effective against the majority. (See Table 3 for details.)

### 4.2. History and virulence spectrum of Stem rust (*Pgt*) races in Ethiopia

The the stem rust race composition in ethiopia were dictated by different factors among the factors the geneics make up of varieties, climate change or weather variations are the major factor that limit

**Figure 1.**
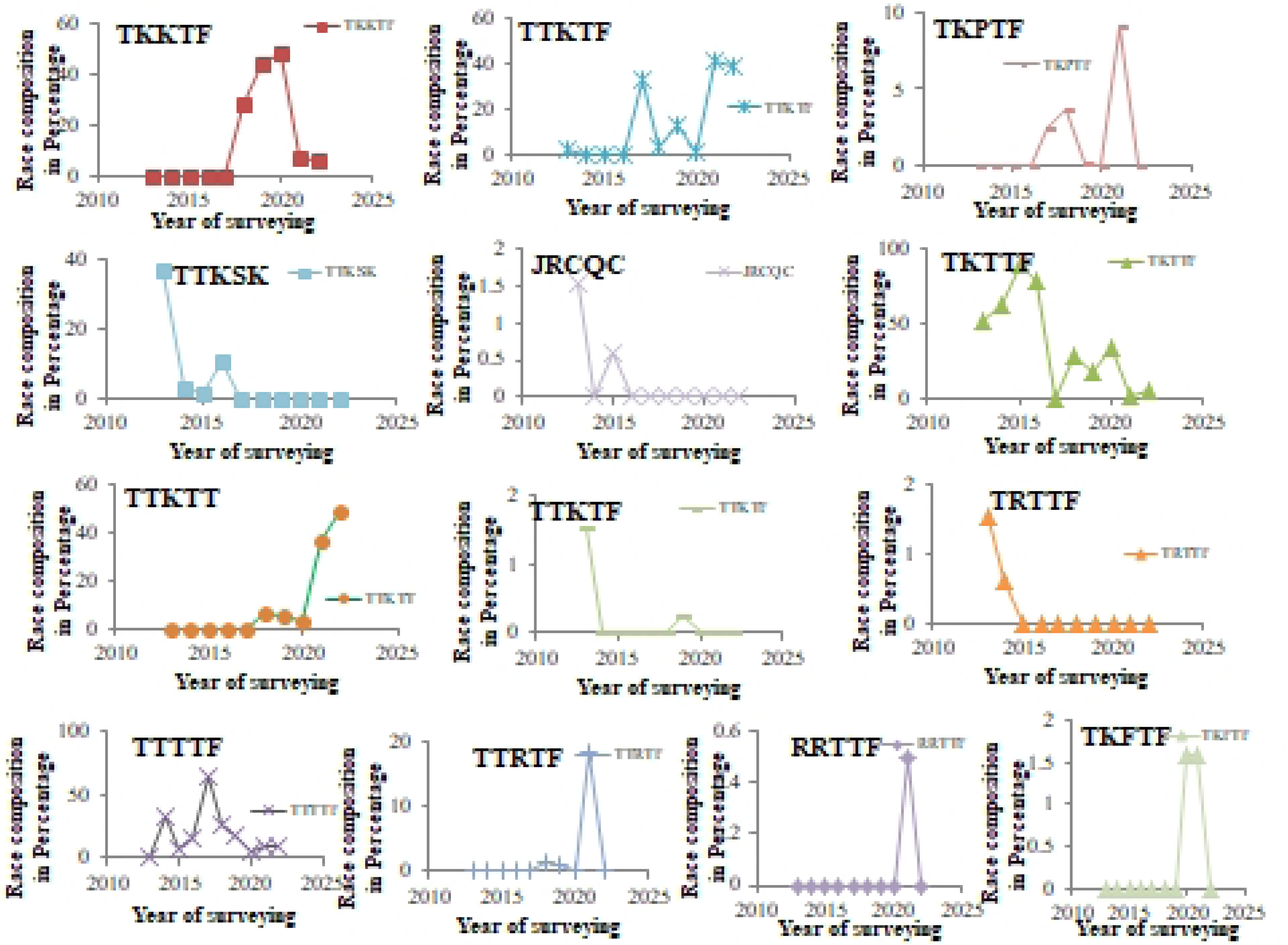
The confirmed wheat rust races and their dynamics in Ethiopia for the decade

Since wheat stem rust is a key yield-limiting disease of wheat in irrigation wheat production areas, this study was designed to identify and examine the re-emergence and extension of recurrently significant crop pathogen races in Ethiopia, *Puccinia graminis* f.sp. *Tritici* races. We identified the predominant stem rust races in Ethiopia’s main wheat-growing regions and considered potential remedies. The second goal of the study was to analyze the racial composition and dynamics of the Ethiopian stem rust population. The study also aimed to assess the resistance levels of bread wheat varieties in relation to the evolving virulent races, identify effective Sr gene sources, and explore the integration of recent scientific advancements for breeding and *Puccinia graminis* f. sp. *tritici* (*Pgt*) management under Ethiopian conditions.

#### 4.2.1. History of stem rust (*Puccinia graminis* f. sp*. Tritici)* races in irrigated wheat production

**Figure 2.**
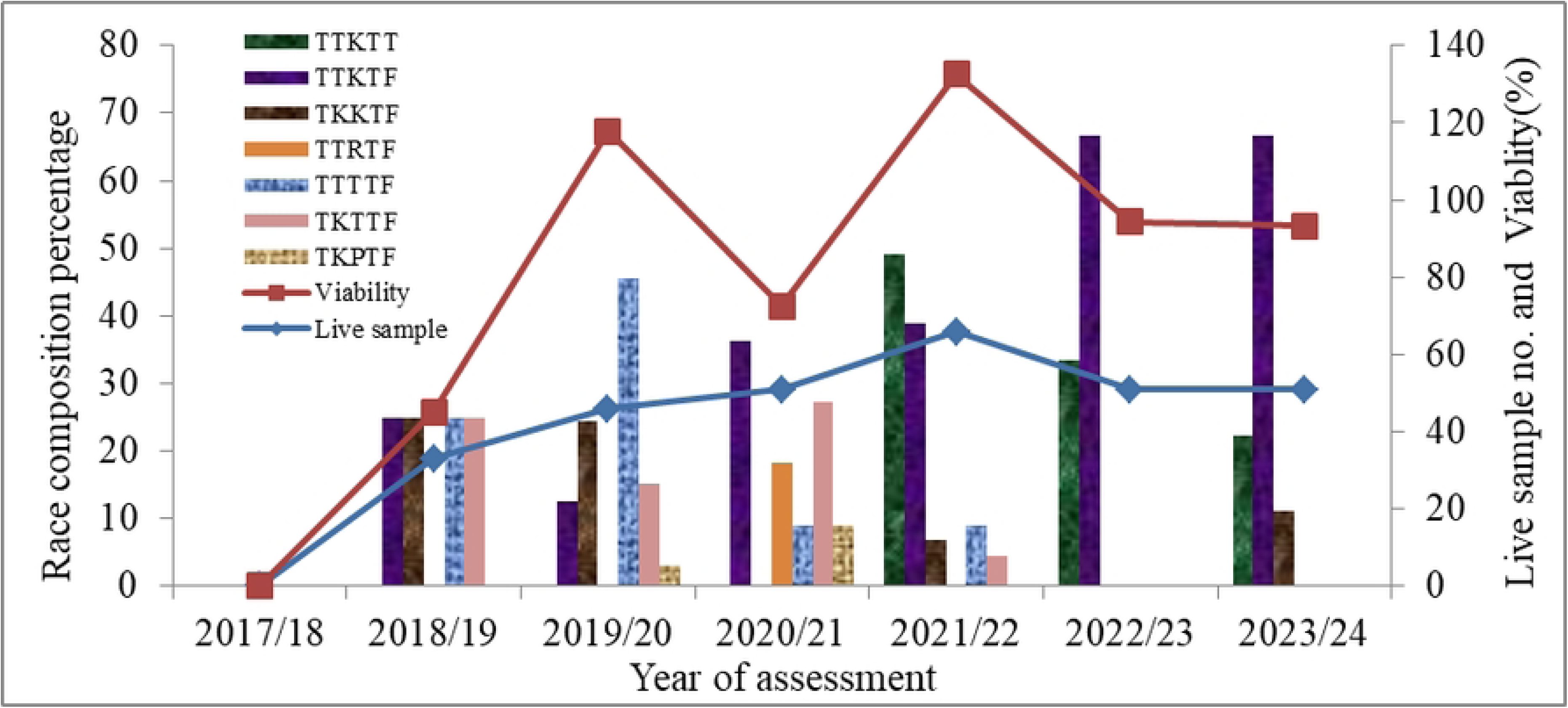
The irrigated wheat rust races and their dynamics in Ethiopia for the decade

The TTKTT race exhibits a 95% virulence spectrum against stem rust resistance genes found in differential lines. Notably, the resistance gene Sr24, which is present in most commercial wheat cultivars worldwide, has lost its effectiveness against this race. Consequently, our breeding programs need to focus on developing more sources of resistance to these aggressive races.

As a result, the eastern region of Ethiopia has become a hotspot for genetically diverse and aggressive races of wheat stem rust (see Table 4). The baseline data provided in this study will be valuable for wheat resistance breeding programs both in this region and globally. Additionally, this data will aid in strategic interventions against stem rust at both national and international levels, supporting efforts to feed the world.

**Table 4.** A virulence-virulence pattern of *Puccinia graminis* f.sp*. tritici* races were identified in the irrigated wheat production areas in the 2021/22 off-season.

| Race | Virulence *Sr* genes | Avirulence *Sr* genes | Virulence frequency (%) |
| --- | --- | --- | --- |
| TTKTT | 5, 6, 7b, 8a, 9a, 9b, 9d, 9e, 9g, 10, 11, 17, 21, 24, 30, 31, 38, *Tmp*, *McN* | 36 | 95 |
| TTKTF | 5, 21, 9e, 7b, 11, 6, 8a, 9g, 9b, 30, 17, 9a, 9d, 10, *Tmp*, 38, *McN* | 36, 24, 31 | 85 |
| TKKTF | 5, 21, 9e, 7b, 6, 8a, 9g, 9b, 30, 17, 9a, 9d, 10, *Tmp*, 38, *McN* | 11, 24, 31, 36 | 80 |
| TTRTF | 5, 21, 9e, 7b, 11, 6, 8a, 9g, 36, 9b, 17, 9a, 9d, 10, *Tmp*, 38, *McN* | 30, 24, 31 | 85 |
| TTTTF | 5, 21, 9e, 7b, 11, 6, 8a, 9g, 36, 9b, 30, 17, 9a, 9d, 10, *Tmp*, 38, *McN* | 24, 31 | 90 |
| TKTTF | 5, 21, 9e, 7b, 6, 8a, 9g, 36, 9b, 30, 17, 9a, 9d, 10, *Tmp*, 38, *McN* | 11, 24, 31 | 85 |
| TKPTF | 5, 21, 9e, 7b, 6, 8a, 9g, 36, 30, 9g, 9a, 9d, 10, *Tmp*, 38, *McN* | 11, 9b, 24, 31 | 80 |

*Sr*, stem rust resistance gene

Irrigated wheat in Ethiopia faces challenges from Puccinia graminis f.sp. Tritici (Pgt), which consists of seven physiological races, including TTKTT and TTKTF. In our research, we identified the races TKKTF, TTRTF, TTTTF, TKTTF, and TKPTF. The most prevalent race, TKKTF, was found in 40 samples (48.2%), followed by TKTTF (the Digelu race) in 28 samples (33.7%). The less common races were TTTTT, TTKTT, and TTTTF, which accounted for one (1.2%), two (2.4%), and four (4.8%) samples, respectively. These races eliminated between 80% and 100% of the resistance genes in various host lines. Notably, the resistance genes Sr24 and Sr31 were effective against the majority of the identified races.

A compilation of several studies (Hailu et al., 2015; Hei et al., 2018; Hailu et al., 2016; Yehizbalem et al., 2020; Hei et al., 2020; Gutu et al., 2021; Gutu et al., 2022; David et al., 2022) indicates that the dynamics of wheat stem rust races are quite high. The data demonstrates that the Ethiopian stem rust pathogen races have evolved significantly, with more virulent strains like TTKTT, TTKTF, and TTTTF becoming dominant. Between the years 2021 and 2022, the aggressive TTKTF race increased its prevalence to 40.64% and 38.8%, respectively.

TKKTF Clade IV-F, which contains a single race (TKKTF), was first discovered in Europe in 2018, although it had previously been identified in Georgia in 2014 and later in Azerbaijan, Iraq, and Eritrea (GRRC) in 2015 and 2016 (Olivera et al., 2019). TTRTF Clade III-B, consisting of one race (TTRTF) and five genetically distinct but related Multi-Locus Genotypes (MLGs), may indicate a resurgence of stem rust at epidemic levels in Europe (Bhattacharya, 2017). The earliest observation dates back to 2014 in Georgia, where it was present at low frequency in a genetically diverse population (Olivera et al., 2019). In the same year, the most prevalent MLG and race within Clade III-B were found in Eritrea and Ethiopia.

According to Hei et al. (2018), TTTTF constituted 32.4% of the viable races in 2014, with over 60% of the isolates from northern Ethiopia, particularly in Gojam and Gondar, being TTTTF (Yehizbalem et al., 2020). The TTTTF race originated from both wheat and Berberis plants (Akci, 2021).

The TKTTF race was first identified in 2012 in southeastern Ethiopia and significantly impacted the popular variety Digelu during the 2013–2014 crop season, causing widespread outbreaks of wheat stem rust (Olivera et al., 2015). With a frequency of 36.4%, TKTTF was the second most common and virulent race (Hailu, 2015). Outbreaks were also reported in Germany in 2013, with TKTTF present in both Clade-A.1 and Clade-A.2 (Olivera Firpo et al., 2017). Within East Africa, Digelu was identified as the most prevalent virulent race and placed in Clade IV-A.1 and Clade IV-B (Olivera et al., 2015).

Recently, TKTTF was confirmed in the United Kingdom and Ireland (Tsushima et al., 2022; Patpour, 2022), likely having transferred from highland regions in Eastern Africa, such as Ethiopia. Clade IV-B, first detected in Croatia in 2017, became widespread in Western Europe between 2019 and 2021. Notably, the TKPTF race was identified only once at Farta in South Gondar in 2017 (Azmeraw et al., 2020), with successful identification occurring in the East Shewa zone’s Ada’a districts in 2019–2020 (Ashagre, 2022). Additionally, these races were observed in regions where irrigation is used for wheat production.

From 2013 to 2017, TTTTF emerged as the most virulent race before being succeeded by TKTTF. Although TKTTF remained common, the distribution and spread of TTKTT are concerning. The results obtained from isolates in the latest studies, particularly at GRRC (Ireland: 2020; UK: 2019, 2021), confirmed the presence of Clade IV-B and race TKTTF in these countries. Historically, this genetic group has been widespread in both East Africa and the Middle East (GRRC), with both the TKTTF and TTTTF races being present in East Africa. The majority of resistance genes were ineffective against most of the races identified in this study. For instance, 13 differential hosts carrying resistance genes 5, 21, 7b, 6, 8a, 9g, 17, 9a, 9d, 10, Tmp, 38, and McN were ineffective against all the identified races in the season (see Table 4). Except for the races TTKTT and TKKTT, only the resistance genes Sr24 and Sr31 were effective against five of the races identified in this study.

Irrigated wheat in Ethiopia faces significant challenges from the pathogen Puccinia graminis f.sp. Tritici (Pgt), which has seven physiological races, including TTKTT and TTKTF. In our study, we identified five races: TKKTF, TTRTF, TTTTF, TKTTF, and TKPTF. The most prevalent race, TKKTF, was found in 40 samples (48.2%), followed by TKTTF (the Digelu race), which was found in 28 samples (33.7%). Other races, TTTTT, TTKTT, and TTTTF, were less common, found in three samples each: one sample (1.2%) for TTTTT, two samples (2.4%) for TTKTT, and four samples (4.8%) for TTTTF. These races effectively eliminated 80 to 100% of the resistance genes in diverse host lines, while the resistance genes Sr24 and Sr31 proved effective against most of the detected races.

Based on a compilation of multiple studies (Hailu et al., 2015; Hei et al., 2018; Hailu et al., 2016; Yehizbalem et al., 2020; Hei et al., 2020; Gutu et al., 2021; Gutu et al., 2022; and David et al., 2022), the dynamics of wheat stem rust races in Ethiopia were notably high. The data indicate that the Ethiopian stem rust pathogen races are highly dynamic, with more virulent strains, such as TTKTT, TTKTF, and TTTTF, emerging and becoming dominant. The TTKTF race, identified in both Berberis and wheat plants (Akci, 2021), was particularly aggressive, increasing its prevalence to 40.64% in 2021 and 38.8% in 2022.

The TKKTF Clade IV-F, which includes the TKKTF race, was first discovered in Europe in 2018, although it had previously been identified in Georgia in 2014 and in Azerbaijan, Iraq, and Eritrea (GRRC) during 2015 and 2016 (Olivera et al., 2019). TTRTF Clade III-B contains one race (TTRTF) and five genetically distinct but related multilocus genotypes (MLGs), signaling a potential resurgence of stem rust at epidemic levels in Europe (Bhattacharya, 2017). Although the initial observation of this clade dates back to 2014 in Georgia, where it was sampled at low frequency within a genetically diverse population (Olivera et al., 2019), the most prevalent MLG and race within Clade III-B were also found in Eritrea and Ethiopia in the same year (Patpour et al., 2020).

According to Hei et al. (2018), 32.4% of the viable races in 2014 were TTTTF. In northern regions of Ethiopia, particularly in Gojam and Gondar, more than 60% of the isolates belonged to the TTTTF race (Yehizbalem et al., 2020). The TTTTF race is believed to have originated from both wheat and Berberis plants (Akci, 2021).

TKTTF was first discovered in 2012 in the southeastern regions of Ethiopia, where it affected the popular wheat variety Digelu during the 2013–2014 crop season, leading to epidemics of wheat stem rust (Olivera et al., 2015). The TKTTF race, specifically the Digelu race, was the second most prevalent and virulent race, with a frequency of 36.4% (Hailu, 2015). Outbreaks occurred in Germany in 2013, and TKTTF was identified in both Clade A.1 and Clade A.2 (Olivera Firpo et al., 2017). Digelu was the most widespread virulent race in East Africa, categorized under Clade IV-A.1 and IV-B (Olivera et al., 2015).

Recently, race TKTTF has been confirmed in the United Kingdom and Ireland (Tsushima et al., 2022, and Patpour, 2022), likely due to its transfer from Eastern African highland regions, such as Ethiopia. Clade IV-B was first detected in Europe in Croatia in 2017 and has become widespread between 2019 and 2021. In 2017, races of TKPTF were recorded solely in a single location in Farta, South Gondar (Azmeraw et al., 2020). However, only the Ada’a districts in the East Shewa zone saw significant success with the TKPTF race during the 2019–2020 season (Ashagre, 2022). Additionally, these races were noted in areas with irrigated wheat production. From 2013 to 2017, TTTTF outperformed TKTTF as the most virulent and aggressive race, but TTKTF later regained prominence. While the TTKTF race remains common, the distribution and expansion of TKTFT are concerning.

Research utilizing isolates from a recent study at the GRRC (Ireland: 2020; UK: 2019, 2021) confirmed the presence of Clade IV-B and race TKTTF in these countries. This genetic group has historically been widespread in both East Africa and the Middle East (GRRC), with East Africa being home to both TKTTF and TTTTF races, indicating a close relationship between them.

The majority of resistance genes proved ineffective against most races identified in this study. For instance, 13 differential hosts carrying resistance genes 5, 21, 7b, 6, 8a, 9g, 17, 9a, 9d, 10, Tmp, 38, and McN were ineffective against all races in the season (Table 4). Except for races TTKTT and TKKTT, only Sr24 and Sr31 were found to be effective against five of the races documented in this study.

Long-distance dispersal of rust races may alter the genetic composition of the rust population (Kolmer, 2005). The races documented in this study exhibited a wide range of virulence. Among the standard differential genes, the dominant race TKKTF was only avirulent to Sr11, Sr36, Sr24, and Sr31, but was virulent against the other genes.

The broadest virulence spectra were observed in the race TTKTT, followed by TTTTF and TKKTT, rendering 95%, 90%, and 90% of Sr genes ineffective, respectively. The races TTKTT, TTTTF, and TKKTT displayed the widest virulence spectrum, affecting 95%, 90%, and 90% of Sr genes, respectively. Most Ethiopian wheat cultivars contain Sr24, to which races TTKTT and TKKTT are virulent. Similarly, in wheat differential lines, race TKTTF was able to overcome 85% of the Sr genes, with only Sr11, Sr24, and Sr31 showing resistance (Table 4).

### 4.3. The genetic race analysis results

#### 4.3.1. The genetic race dynamics analysis results in wheat production areas of Ethiopia

Clade II-B was the most dominant race throughout the years 2014 to 2017, according to the results of the physiological race analysis, which was also supported by the genetic race analysis results. TKTTF, followed by TKKTF and TTRTF, was the most important race from 2014 through 2022 (Figure 3). The dynamics also supported the shift in the virulent race’s frequency of occurrence in the years 2021 and 2024, when clade IV-F completely dominated the clade distribution (Figure 4).

**Figure 3.**
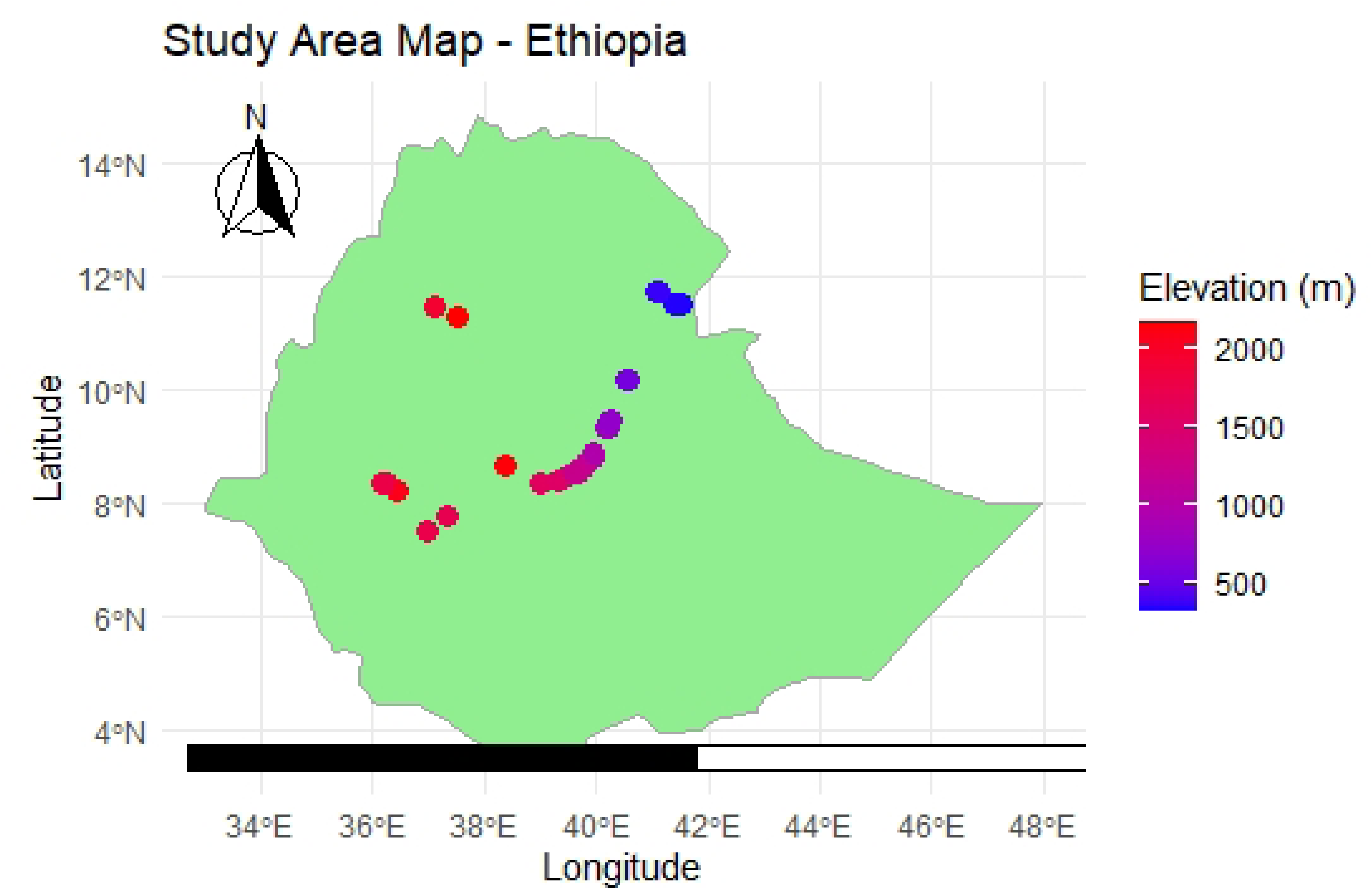
The wheat rust distribution map with elevation model dynamics in Ethiopia for the past decade period

**Figure 4.**
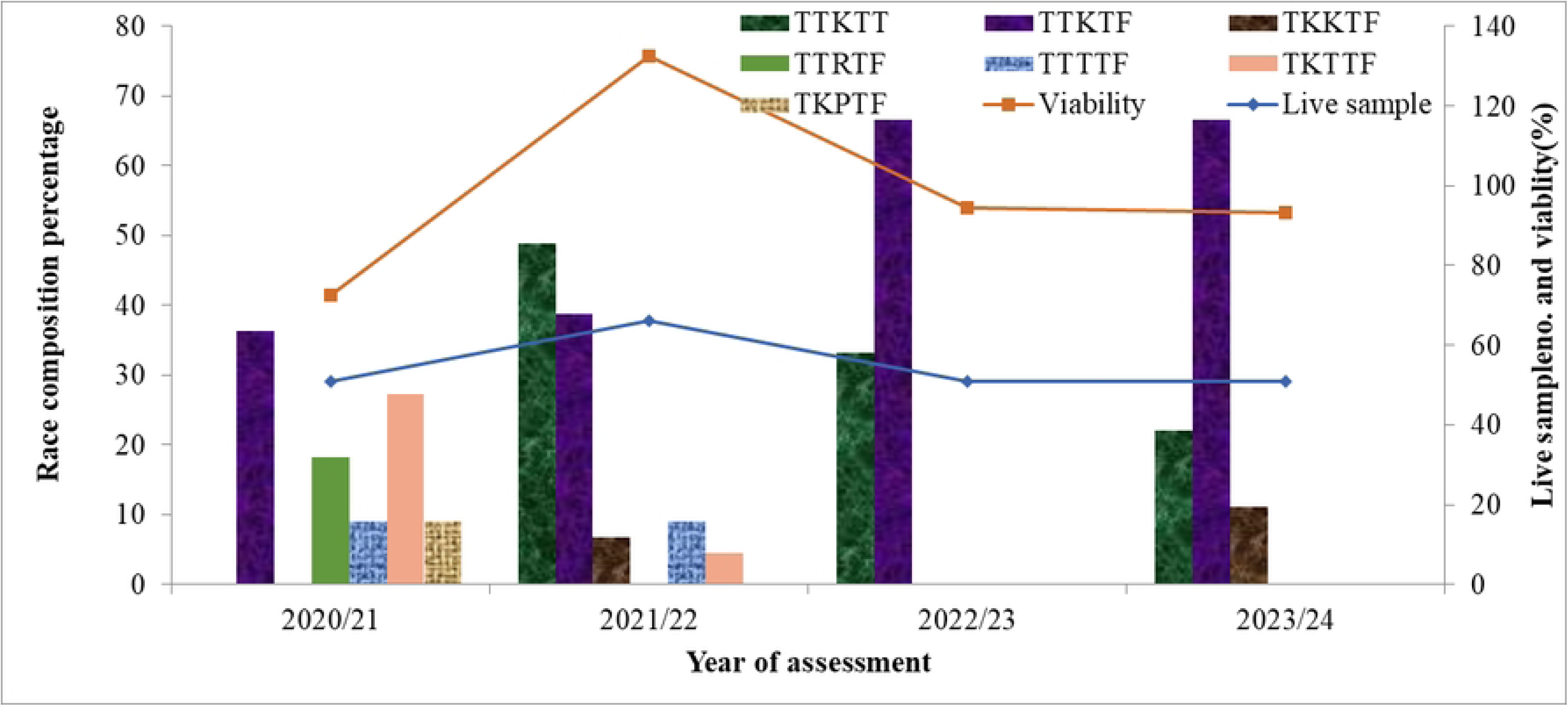
The wheat rust distribution race map with elevation model dynamics in Ethiopia for the past decade period

##### Step 2: Heat Map Generation

Heat Map Generation

The heat map above illustrates the intensity of various rust races across Ethiopia. According to the data, the TTKTF race is the most common, appearing frequently across multiple years and locations, particularly in central and eastern Ethiopia. The TTKTT race was also noted in 2021/22 and 2022/23, primarily concentrated in higher elevation areas.

In contrast, the races TTRTF and TTTTF were less common but were observed in specific regions, such as the northern and central parts of the country. TKKTF was the dominant race found in lower-elevation areas. The TKTTF and TKPTF races had sporadic occurrences, mainly in lower-elevation regions.

In 2020/21, there was a high intensity of TTKTF in central Ethiopia (latitude ∼9.34, longitude ∼40.18). In 2021/22, TTKTT spread in higher elevations (latitude ∼8.34, longitude ∼38.98). By 2022/23, TTKTF continued to be present in central regions, while TTKTT appeared in southern areas. In 2023/24, TTKTF emerged in northern Ethiopia (latitude ∼11.73, longitude ∼41.09).

As indicated in the four-year temporal distribution figure (Figure 4), the two main races in irrigated areas were TTKTT and TTKTF.

##### Distribution Maps

TTKTF is primarily concentrated in central Ethiopia, with some expansion to the north and south over the years. TTKTT is mainly found in higher elevation areas, exhibiting a noticeable increase in 2021/22. In contrast, TTRTF and TTTTF are limited to specific regions and have not shown any significant spread over the years (Fig. 5).

**Figure 5.**
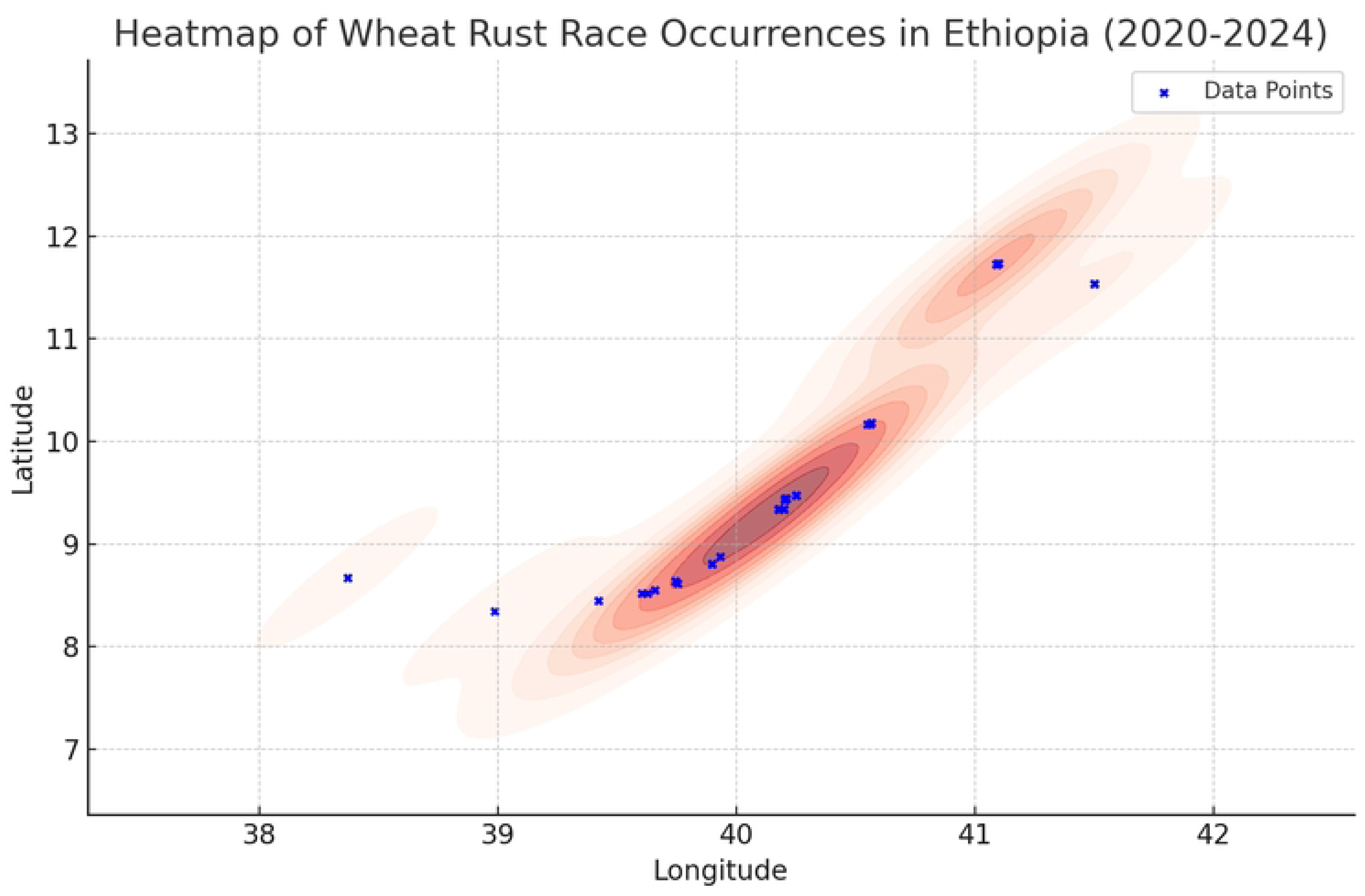
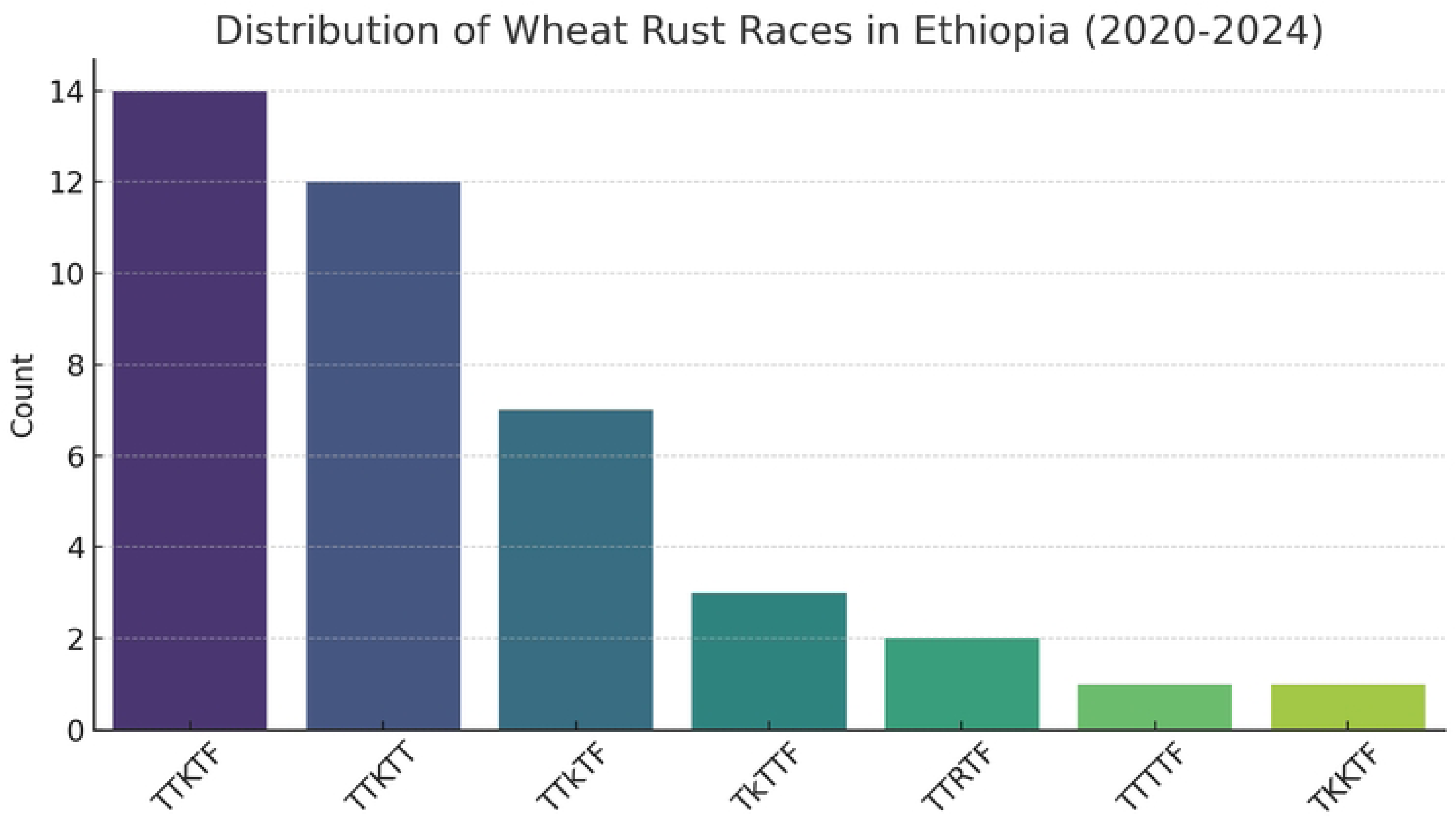
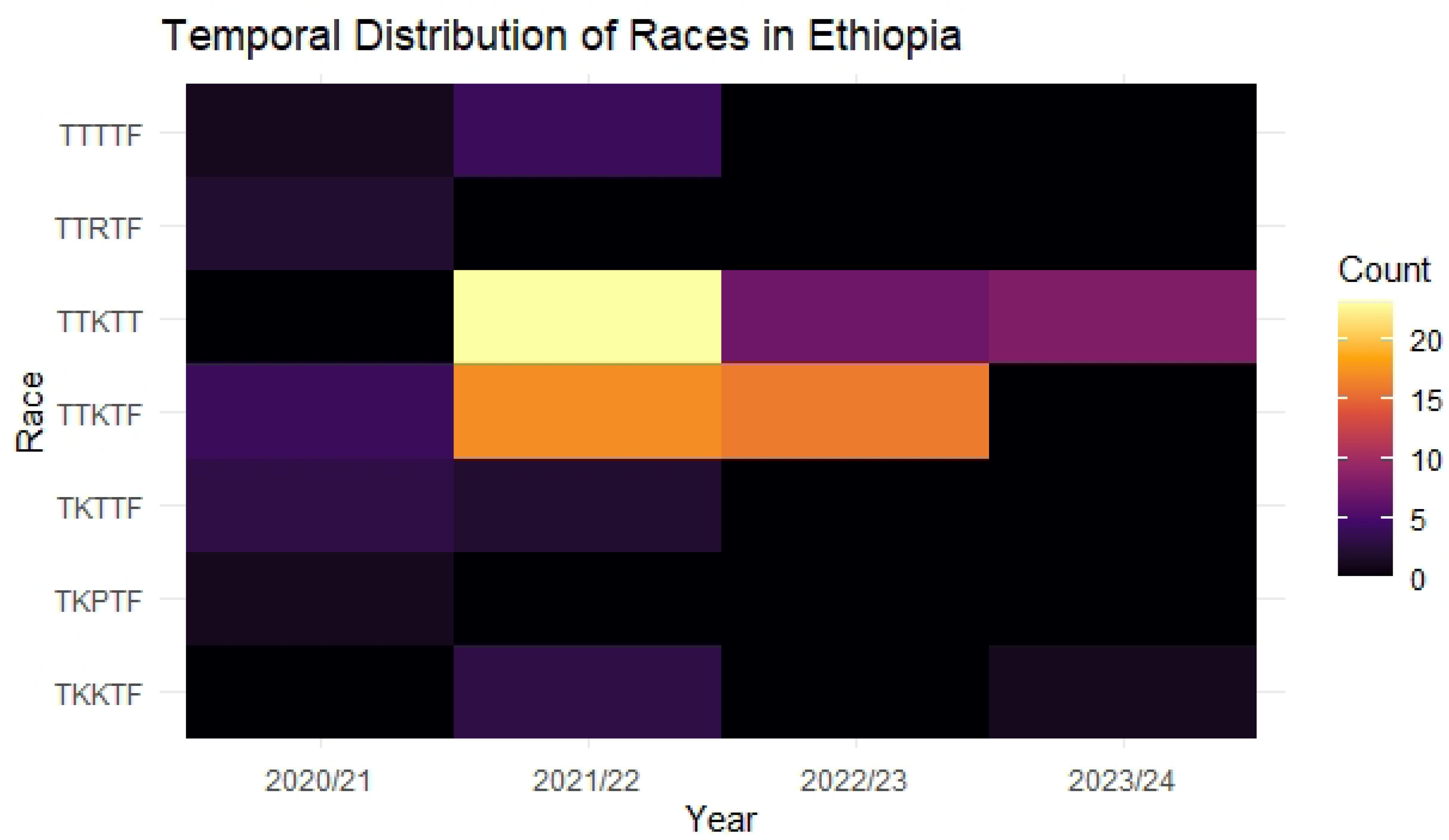
The wheat rust physiological races dynamics heat composition in irrigated wheat production areas of Ethiopia for the four years

#### 4.3.2. The genetic race analysis results in irrigated wheat production areas of Ethiopia

In 2019, nine samples from the Minnesota CDL lab were genotyped as belonging to clade III-B (Co-A04), which is associated with the Pgt race TTRTF. According to these data, the Pgt races TTRTF (III-B) and TKFTF/TKKTF (IV-F) have emerged as the dominant races during this period, which aligns with race phenotyping research conducted in Ambo.

### 4.4. Distribution of Pgt races in the irrigated wheat production areas of Ethiopia

The wheat varieties Amibara, Kakaba, and an unknown variety were reported to be infested by the race TKKTF in the West Arsi, East Shoa, and Jimma regions. In West Arsi and East Shoa, the race TKTTF was found infesting the wheat varieties Fentale 2 and Ogolcho. In West Arsi, the race TTTTF was discovered infesting an unknown wheat variety known as Limmu.

Furthermore, in West Arsi, East Shoa, Buno Bedele, Jimma, and Awi, the wheat varieties Amibara, Kakaba, Ogolcho, Kingbird, Wane, and Danda’a were found to be affected by the race TTKTF. The varieties Kakaba, Danda’a, and Oborra were identified as being impacted by the race TTKTT in Jimma and Buno Bedele.

Additional findings included variants such as TTTTF affecting Denda’a, Kakaba, Digelu, and an unknown variety called Honqoltu, along with other cultivars like Lakech, TTKTT, ETBW 9553, and Ogolcho. The stem rust survey also indicated that the majority of widely planted wheat cultivars were impacted by stem rust races, including Ogolcho, Kubsa, Hidase, Lakech, Digelu, Denda’a, ETBW 9553, Lemu, Kingbird, and Wane, as well as some unidentified wheat cultivars. (Table 5 summarizes the affected cultivars and their corresponding races.)

**Table 5.**
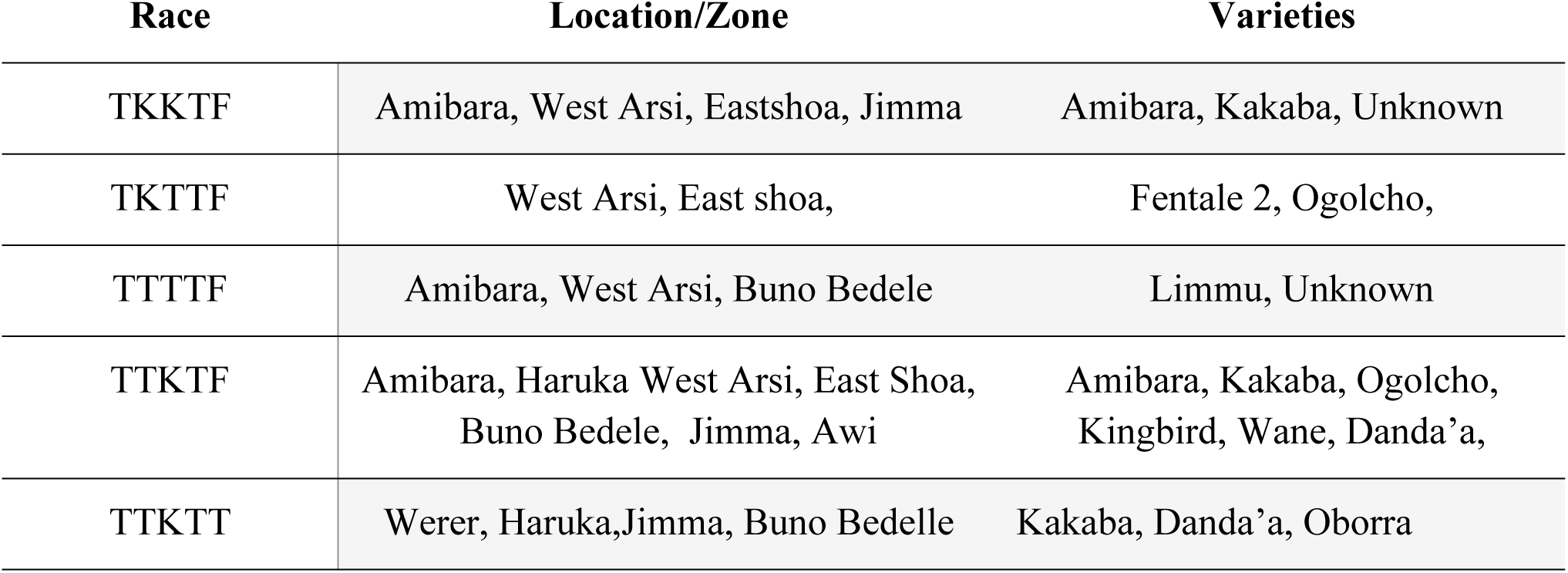
Locations and varieties from which the races were identified from irrigated areas in 2021/22-2023/24.

**Table 6.** Virulence spectrum of the *Pgt* races identified during the main season of 2018-2022 in irrigated wheat production areas in Ethiopia.

| Races | Virulence | Avirulence |
| --- | --- | --- |
| TTTTF | 5, 21, 9e, 7b, 11, 6, 8a, 9g, 36, 9b, 30, 17, 9a, 9d, 10, Tmp, 38, McN | 24, 31, |
| TKTTF | 5, 21, 9e, 7b, 6, 8a, 9g, 36, 9b, 30, 17, 9a, 9d, 10, Tmp, 38, McN | 11, 24, 31 |
| TKKTF | 5, 21, 7b, 6, 8a, 9g, 9b, 30, 17, 9a, 9d, 10, Tmp, 9e, 38, McN | 11, 36, 24, 31 |
| TTRTF | 5, 21, 9e, 7b, 11, 6, 8a, 9g, 36, 9b, 17, 9a, 9d, 10, Tmp, 38, McN | 30, 24, 31 |
| TTKTF | 5, 21, 9e, 7b, 11, 6, 8a, 9g, 9b, 30, 17, 9a, 9d, 10, Tmp, 38, McN | 36, 24, 31 |
| TTKTT | 5, 21, 9e, 7b, 11, 6, 8a, 9g, 9b, 30, 17, 9a, 9d, 10, Tmp, 24, 31, 38, McN | 36 |
| TKPTF | 5, 21, 9e, 7b, 6, 8a, 9g, 30, 17, 9a, 9d, 10, Tmp, 38, McN | 11, 36, 9b, 24, 31 |

### 4.5. Altitudinal Distribution of Pgt races in the Irrigated Wheat Production areas of Ethiopia (2019-2024)

TKKTF was identified in West Arsi, East Shoa, and Jimma, specifically on the wheat variety Amibara I, in the rain-fed agroecologies of Ethiopia. The primary races were found in lowland and desert regions where wheat is cultivated. The wheat variety Fentale 2 was particularly infested by the races TKTTF (77.73%) and TKKTF (60%). The most virulent strains demonstrated a broader ecological adaptability, thriving in environments ranging from lowland deserts to the highest altitudes. Additionally, TTTTF and TTKTF were detected in midland, lowland, and desert plain agroecologies at rates of 11.1%, 52.96%, and 35.93%, respectively.

**Table 7.**
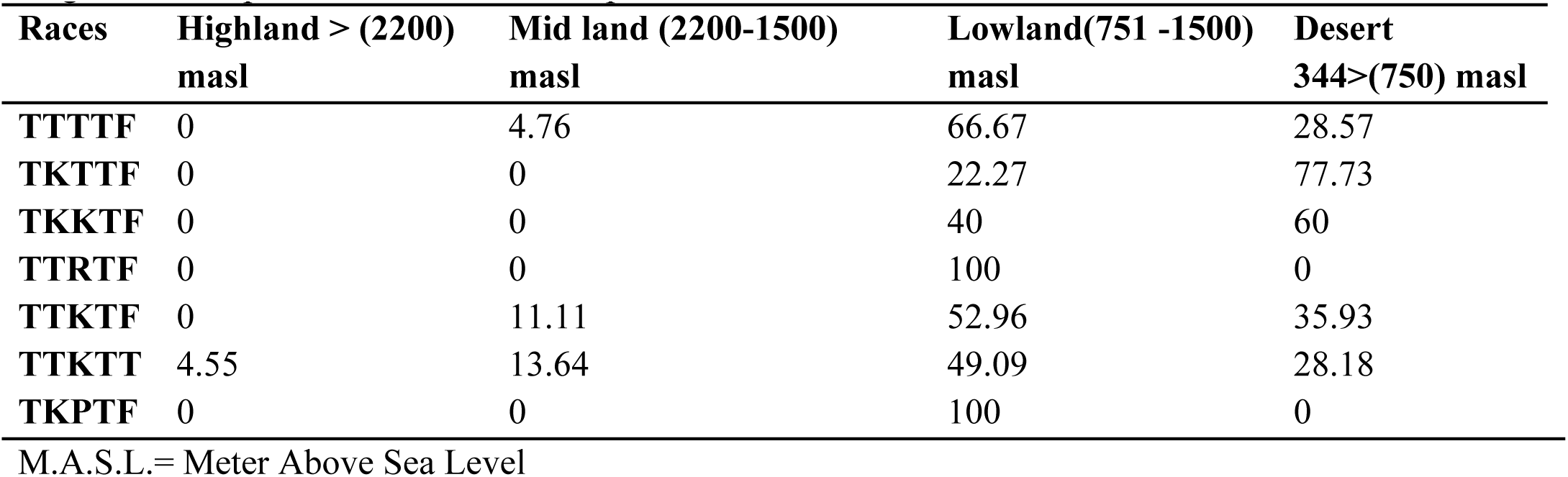
Pgt races distribution identified during irrigated wheat main season of 2021-2023/24 on irrigated wheat production areas in Ethiopia.

| Races | Highland > (2200)<br>masl | Mid land (2200-1500)<br>masl | Lowland(751 -1500)<br>masl | Desert<br>344>(750) masl |
| --- | --- | --- | --- | --- |
| TTTTF | 0 | 4.76 | 66.67 | 28.57 |
| TKTTF | 0 | 0 | 22.27 | 77.73 |
| TKKTF | 0 | 0 | 40 | 60 |
| TTRTF | 0 | 0 | 100 | 0 |
| TTKTF | 0 | 11.11 | 52.96 | 35.93 |
| TTKTT | 4.55 | 13.64 | 49.09 | 28.18 |
| TKPTF | 0 | 0 | 100 | 0 |

M.A.S.L.= Meter Above Sea Level

### 4.6. Climate Change in Ethiopia and Implications for Wheat Stem Rust (Pgt)

Ethiopia is known as one of the most climate-vulnerable countries in sub-Saharan Africa. Its location, dependence on rain-fed agriculture, and reliance on smallholder farming make it especially sensitive to climate changes. Evidence from observations and projections shows that the country has already faced significant climate shifts over the last fifty years likely it faces similar trends for the coming 10 years.

#### Observed Trends and Near-Term Projections (to 2030)

From 1960 to 2006, Ethiopia’s average annual temperature rose by about 1.3 °C, which equals a warming rate of 0.28 °C per decade. This warming has been accompanied by an increase in extreme heat events in several regions, especially in the Upper Blue Nile Basin and the northeastern highlands. At the same time, rainfall patterns have become more unpredictable. Some areas, like parts of Wollo, have seen slight increases in Kiremt (June–August) rainfall, while significant declines have occurred in Belg (March–May) and Bega (December–February) rains. These changes disrupt traditional cropping calendars and increase year-to-year variability, creating serious challenges for smallholder wheat production (Simane *et al*., 2016; Mohammed et al., 2022; Mohammed et al., 2025; Adane & Asmerom, 2025).

#### Near-Term Projections (to 2030)

Projections from CMIP6 and CORDEX-Africa ensembles suggest that warming will continue across Ethiopia, regardless of the emission scenario. Under SSP1-2.6, average annual temperatures are expected to rise by about 0.5–0.7 °C above the 1995–2014 average by 2030. Under SSP2-4.5, this increase could reach around 0.8–1.0 °C, while SSP5-8.5 predicts a rise of about 1.2 °C in that same period. Rainfall projections show greater uncertainty due to Ethiopia’s varied topography and climate. However, average models suggest slight declines under low- and moderate-emission scenarios (SSP1-2.6, SSP2-4.5). Some models under high-emission scenarios (SSP5-8.5) predict slight increases in annual rainfall, along with more extreme weather events (Rettie et al., 2023; IPCC, 2021).

**Fig 6.**
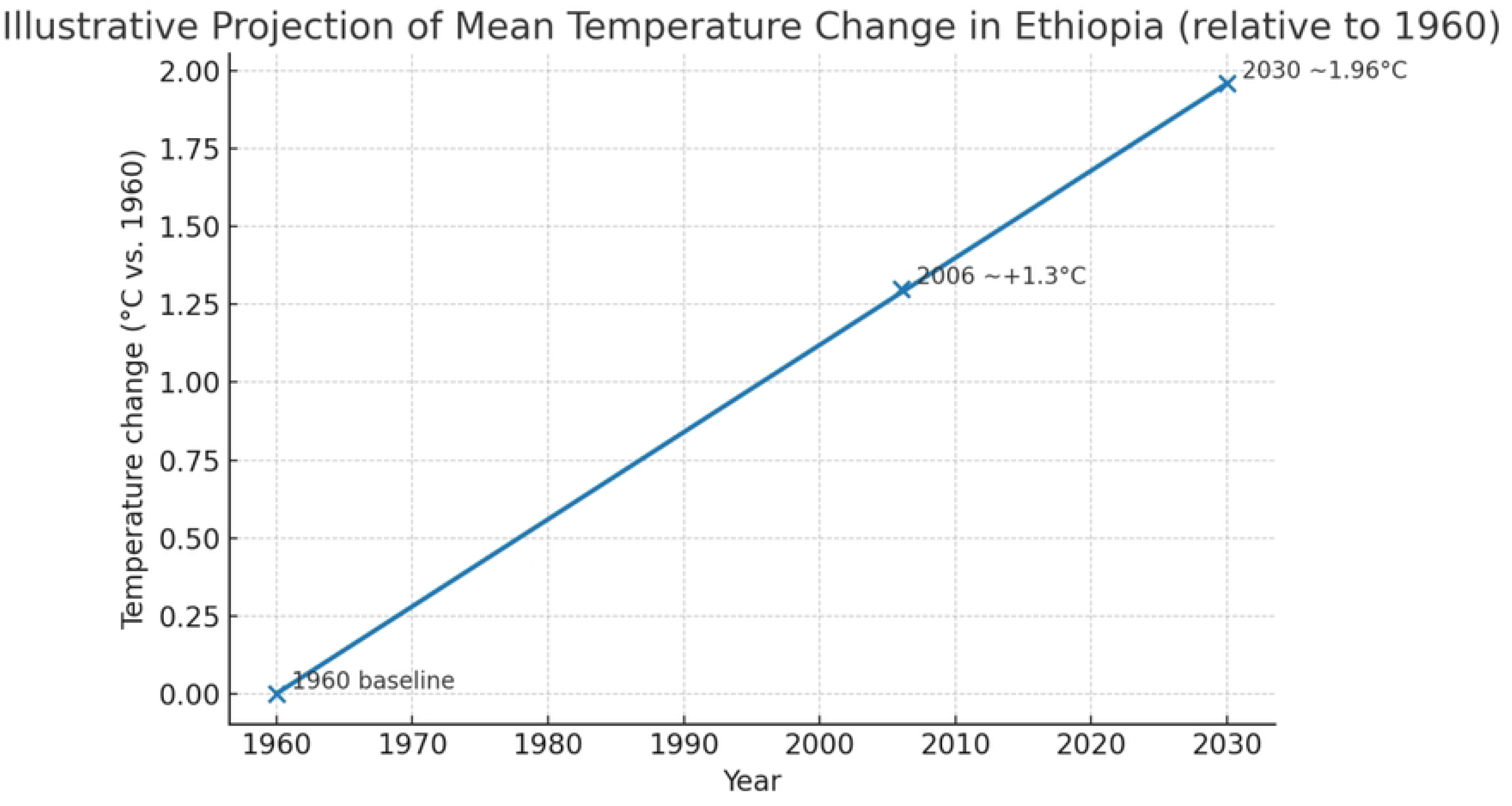
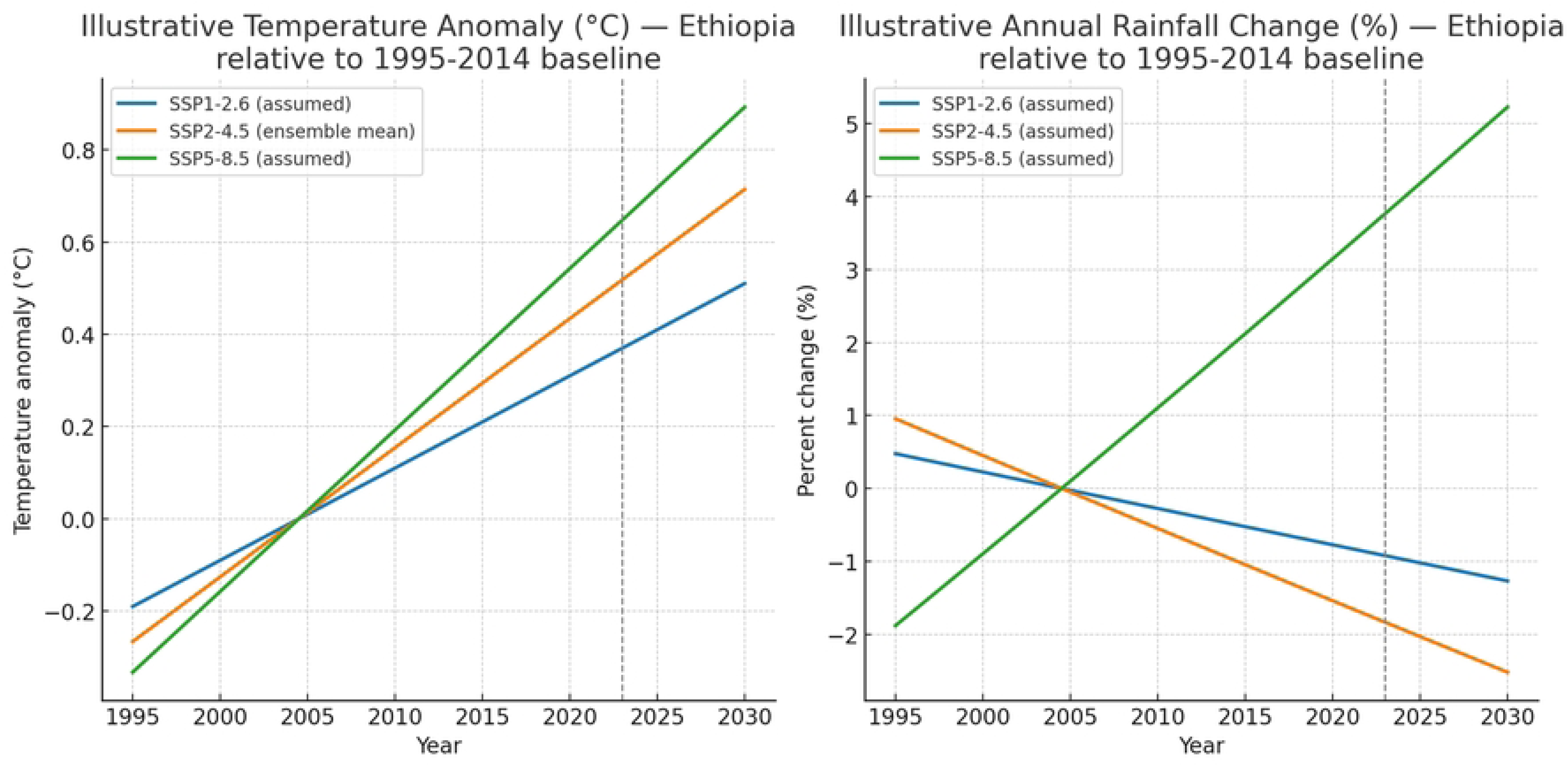
Trend of climate change in Ethiopia a. mean temperature change b. anomaly and temperature change and annual rain fall change based up on 1995-2014.

#### 4.6.2. Implications for Wheat Stem Rust

Warmer conditions create more favorable environments for the proliferation of pests and pathogens. For instance, wheat stem rust (*Puccinia graminis* f. sp. *tritici*) is projected to expand into previously cooler highland areas as minimum night temperatures rise. The increase in stem rust epidemics in Ethiopia is closely tied to these climate changes. Stem rust (*Puccinia graminis* f. sp. *tritici*) flourishes in warm and humid conditions, especially where night temperatures remain moderate and leaf wetness lasts long enough for spore germination. Warmer minimum temperatures, projected to rise faster than daily maximums, will expand the areas where rust can thrive, allowing the pathogen to move into regions that were previously less affected.

From the above premises we are defiantly in climate change. The evidence shows that Ethiopia faces a twofold challenge: rising temperatures and increasingly unpredictable rainfall patterns will change the regions suitable for wheat production and increase the threat of stem rust outbreaks. By 2030, under all likely SSP scenarios, the warming trend will strongly influence pathogen behavior. In this context, it is crucial to combine climate projection data with plant disease models for better breeding and early-warning systems. Ethiopia, due to its key role in wheat production and genetic diversity, is at a crucial point for ensuring both national food security and global agricultural stability.

The interaction between climate change and *Puccinia graminis* f. sp. tritici (Pgt) is not only about increasing incidence, but also about the evolution, distribution, and epidemiology of new and existing races. Impact of Climate Change on new and existing *Puccinia graminis* f. sp. *tritici* (*Pgt*) races are expected in multi facet modes. These includes Acceleration of pathogen evolution, Geographic expansion of existing races, Increased survival and overwintering of inoculum, Emergence of highly virulent races and will challenges the resistance breeding development.

Acceleration of pathogen evolution will rise temperatures and longer periods of suitable climate increase the number of infection cycles that *Pgt* can complete in a single growing season. This increase in generation turnover creates more chances for mutation and genetic recombination, raising the likelihood of new virulent races appearing. Research shows that larger pathogen populations and more frequent epidemics speed up evolutionary changes in plant pathogens (Singh et al., 2023; Arenas et al., 2018). Climate-related stress in wheat, such as from heat or drought often reduces the effectiveness of quantitative and some major-gene or R resistances. This allows greater pathogen reproduction on partially susceptible hosts. Such stress can act as a filter, favoring virulent variants that were previously rare (Velásquez, 2018; Mao *et al*., 2023).

Geographic expansion of existing stem rust races with global warming trends are expected to make higher altitudes and cooler agricultural zones more suitable for stem rust. In Ethiopia, this means that highland wheat areas (around > 2,000 m.a.s.l.) may experience conditions that support *Pgt* life cycles. As a result, aggressive lineages, including the Ug99 group and its derivatives, may thrive in areas where climate once limited their survival. Field and modeling studies from East Africa show that changes in temperature and precipitation can lead to shifts in the range of rust pathogens (Li *et al*., 2019; Fetch *et al*., 2021). With multiple races existing in a single region, the chances of mixed infections on individual hosts increase. This co-occurrence promotes genetic exchange, whether through sexual recombination or somatic processes, further diversifying the pathogen population (Olivera *et al*., 2019; Arenas *et al*., 2018).

Increased survival and overwintering of inoculum warmer minimum (night-time) temperatures extend the life of airborne urediniospores and increase the time that inoculum can survive between cropping cycles, known as the Ethiopia produces three times a year may serve as “green-bridge.” Studies have shown that urediniospores can remain infectious for long periods at low but non-freezing temperatures. Thus, warmer winters support a larger reservoir of overwintering inoculum. In Ethiopia’s bimodal rainfall systems (Belg and Meher), this extended inoculum window may lead to greater infection carryover between seasons (Singh *et al*., 2023; Barua *et al*., 2018). Changes in the distribution and life cycles of alternate hosts, especially Berberis spp., with climate change will allows their expansion or improved survival, could further increase opportunities for sexual recombination and the creation of new genotypes. Observations of sexual populations and the role of alternate hosts in *Pgt* diversity highlight the epidemiological importance of these changes (Olivera *et al*., 2019).

The emergence of highly virulent races, Ethiopia’s diverse agricultural conditions and its history as a center for *Pgt* diversity create a setting where increased disease pressure, caused by climate change, can speed up the evolution of highly virulent races. More disease and increased fungicide use in favorable climatic conditions may strongly select for fungicide resistance and virulence against resistance genes. Historical and molecular studies of Ug99 and other emerging lineages show how regional selection pressures can lead to the development of highly virulent races with global impacts (Li *et al*., 2019; Fetch *et al*., 2021).

Challenges for resistance breeding resistance genes vary in their stability across environments. Some Sr genes show temperature-sensitive traits, while quantitative (polygenic) resistance is often more effective across different conditions than single, major-gene resistance. Therefore, the genetic improvements made in current climates may not last under future heat stress. The climate-driven emergence of new *Pgt* races calls for breeding strategies that focus on combining multiple complementary resistance genes and incorporating durable, quantitative resistance along with quick monitoring to detect changes in virulence (Gao *et al*., 2019; Mapuranga *et al*., 2022).

Additionally, changes in rainfall patterns may lead to a dual risk. Excessive rainfall and high humidity can create ideal conditions for spore growth and infection. Conversely, drought can weaken the host plants, making wheat more vulnerable when better conditions return. The expected increase in extreme rainfall under SSP5-8.5 could further raise the chances of epidemic years, as strong winds and storms help spread spores.

Ethiopia’s role as a global center of wheat diversity makes these findings even more important. While breeding for resistance has provided some temporary solutions especially against the highly virulent Ug99 race climate-related changes in how pathogens behave may hasten the breakdown of resistance genes. Without strategies like using multiple resistant wheat varieties, fungicide applications, and disease forecasting based on climate data, Ethiopia may face ongoing yield losses that threaten both national food security and the global wheat supply chain (Singh *et al*., 2015; Tesfaye *et al*., 2021; Wubaye *et al*., 2023).

## 5. Conclusion

The past few decades of physiological and genetic analysis of stem rust in Ethiopia’s wheat production areas have revealed frequent race mutations in the race composition. A four-year analysis conducted in the irrigated wheat production regions identified seven distinct races, with a notable shift towards highly virulent forms. The study confirmed a high spectrum of virulence and significant population variability among the seven detected stem rust races. These races are critical because they significantly impact the rain-fed wheat production in the country.

During the 2021/22 period, the most virulent race, TTKTT, was predominant. The identified races included TTKTT (49%), TTKTF (38.8%), TTTTF (6.8%), and TKTTF (4.5%). The results also indicated that the emergence of new virulent races, such as TTKTT, rendered 19 of the 20 tested stem rust differential resistance genes ineffective (95% of the genes), while TTKTF affected 85% of them. Race TTKTT is expanding geographically and many commonly grown wheat cultivars are now susceptible to it. Notably, TTKTT, which was newly identified and most virulent, appeared for the first time in Ethiopia at a frequency of 49%. In contrast, the second most virulent race, TTTTF, saw a decline in frequency from 2019/20 through the subsequent two years, causing 90% of the differential resistance genes to be ineffective.

Race TTKTF was first detected in Ethiopia and exhibits a high virulence spectrum, making 17 of the 20 tested stem rust differential resistance genes ineffective (85%). The dynamics of stem rust races in Ethiopia indicate a shift beginning with TKTTF in 2013, followed by TTTTF in 2017, which was later replaced by TTKTF in 2020. Currently, the two most frequent virulent races are TTKTT and TTKTF. These findings highlight the urgent need for resistance breeding and the exploration of new strategies to combat these persistent races.

The only effective resistance gene identified in commercial wheat varieties, such as Danda’a, is Sr24, which has shown ineffective protection against these races (Kittessa et al., 2021). Therefore, the lack of stem rust resistance in widely cultivated wheat varieties underscores the necessity to initiate new breeding efforts focused on stem rust resistance in ongoing and future breeding programs. These efforts could involve both conventional and novel breeding methodologies (Singh et al., 2015; Rimsha, 2021; Luo et al., 2021; Wulff and Krattinger, 2022). Coordinated action is essential, as the globe, much like a community, must address the introduction of virulent races stemming from mutation hotspots (Patpour et al., 2022).

It is crucial to emphasize the importance of regular, coordinated pathogen and disease surveillance efforts at both national and global levels to avert potential devastation. The Eastern African highlands, known as the Horn of Africa highlands, were significant sources of disease epidemics in Europe and the Middle East. Epidemiological studies comparing the aggressiveness of prevalent clades, races, and their temperature requirements would prove invaluable.

Consequently, breeding programs should prioritize the search for more sources of resistance to virulent races of *Puccinia graminis tritici* (*Pgt*). In Ethiopia, the increasing virulence spectrum of stem rust races threatens the effectiveness of the resistance genes currently in use. Irrigated wheat production merits attention to address potential green bridges and the risk of genetically diverse and virulent races affecting both the belg and main-season crops.

The baseline information generated by this study is crucial for strategic interventions related to rust management, screening, and resistance breeding efforts for wheat breeders and seed technology multiplication units in the region.

The rapid spread of highly virulent stem rust races, particularly TTKTT, underlines the necessity for durable resistance strategies within Ethiopia’s wheat production systems. Incorporating transgene cassettes containing multiple resistance genes into elite wheat cultivars represents a promising long-term disease control strategy. Furthermore, breeding programs must focus on identifying and deploying novel resistance sources to counteract the continuous evolution of Pgt races. Special attention should be paid to irrigated wheat production, as it serves as a critical inoculum reservoir that threatens both belg and main-season wheat crops. Insights from this study provide essential guidance for strategic rust management, resistance breeding, and seed technology development in Ethiopia.

## 6. Recommendation

Targeted monitoring focuses on regions with high rust race intensity, such as central Ethiopia, where TTKTF is prevalent. Elevation-based strategies should be developed to create rust-resistant wheat varieties suited for high-altitude areas, particularly where TTKTT is common. Continuous yearly analysis should be conducted to monitor rust race distribution and identify emerging threats.

## 7. Acknowledgements

We would like to express our sincere gratitude to the “Improved Disease Monitoring and Management for Wheat and Cassava through Epidemiological Modeling” project for its financial support. We are particularly thankful to the Ambo, Deber Markos, Kullumssa, and Werer Agricultural Research Centers of the Ethiopian Institute of Agricultural Research (EIAR), as well as CIMMYT Ethiopia and the Adet Agricultural Research Center (ARARI), for their logistical assistance in conducting the survey. We extend our special thanks to the Center for Disease Learning (CDL) at the University of Minnesota for their valuable genetic race analysis of the Pgt.

## Annex 1. Summary of the paperwork

Study in the climate investigates the race dynamics of wheat stem rust in Ethiopia’s irrigated and rain-fed wheat-producing areas during both the fall and summer seasons, utilizing both primary and secondary data. The findings aim to guide the direction and steps needed for research and development focusing on sustainable wheat production in the country. Achieving sustainability in Ethiopia hinges on overcoming the challenges posed by two economically significant rust pathogens: stem rust (*Puccinia graminis* f. sp. tritici) and yellow rust (*Puccinia striiformis* f. sp. *tritici*).Special attention needed for the *Pgt* because it will expand to new major wheat producing areas.

The research emphasizes the technical foundation and strategies required to provide long-lasting solutions to issues related to recurring infections and race modifications of these pathogens. A comprehensive review of available literature was conducted to better understand the race dynamics of rust pathogens in Ethiopia, focusing on recent events that have posed challenges.

For secondary data, the study referenced various published sources on physiological race analysis, along with annual early warning and monitoring reports by Minnesota_CDL, as well as findings from molecular or genetic analyses conducted at Aarhus University. Following the significant disruptions to global wheat production caused by the UG99 race and its recurring outbreaks in the 2000s, the analysis of wheat rust races has gained international attention.

The main focus of the paper is on the changes in stem rust races in Ethiopia. It analyzes the race history of stem rust (*Puccinia graminis* f. sp. *tritici*), the emergence of new virulent races, and their impact on both adapted and released irrigated wheat varieties in Ethiopia. The study aims to provide a pathway forward based on lessons learned from the 2021 to 2024 production years.

## References

Adane, A., & Asmerom, B. (2025). Analysis of spatiotemporal distribution, variability, and trends of rainfall in Wollo area, Northeastern Ethiopia. PloS one, 20(1), e0312889.

Adimassu, Z., Tamene, L. D., Tibebe, D., Ebrahim, M., & Abera, W. (2023). Identification and prioritization of context-specific climate-smart agricultural practices in major agro-ecological zones of Ethiopia.

Admassu B, Lind V, Friedt W, and Ordon F. 2009. Virulence analysis of *Puccinia graminis* f. sp. *tritici* populations in Ethiopia with special consideration of Ug99. Plant Pathology 58(2):362–369.

Admassu BH, Friedt W, and Ordon F. 2012. Stem rust seedling resistance genes in Ethiopian wheat cultivars and breeding lines. African Crop Science Journal, 20(3): 149–162.

Akci, N., Karakaya, A. *Puccinia graminis* f. sp. *tritici* races identified on wheat and Berberis spp. in northern Turkey. Indian Phytopathology 74, 1105–1109 (2021). 10.1007/s42360-021-00343-1

Arenas, M., Araujo, N. M., Branco, C., Castelhano, N., Castro-Nallar, E., & Pérez-Losada, M. (2018). Mutation and recombination in pathogen evolution: Relevance, methods and controversies. Infection, Genetics and Evolution, 63, 295–306.

Asadnabizadeh, M. (2023). Critical findings of the sixth assessment report (AR6) of working Group I of the intergovernmental panel on climate change (IPCC) for global climate change policymaking a summary for policymakers (SPM) analysis. International Journal of Climate Change Strategies and Management, 15(5), 652–670.

Ashagre ZA (2022) Detection of Wheat Stem Rust (*Puccinia graminis* f.sp. *tritici*) Physiological Races from Major Wheat Producing Regions of Ethiopia. Aquac Fish Stud Volume 4(3): 1–6.

Barua, P., You, M. P., Bayliss, K. L., Lanoiselet, V., & Barbetti, M. J. (2018). Extended survival of Puccinia graminis f. sp. tritici urediniospores: implications for biosecurity and on-farm management. Plant Pathology, 67(4), 799–809.

CSA (Central Statistics Authority). 2019. Report on area and production of major grain crops. Addis Ababa, Ethiopia. 24

Denbel W, Badebo A, and Alemu T. 2013. Evaluation of Ethiopian commercial wheat cultivars for resistance to stem rust of wheat race ‘UG99’. International Journal of Agronomy and Plant Production, 4(1): 15–24.

Eshetu B., (1985) Review of research on diseases of barley, tef, and wheat in Ethiopia, A review of crop protection research in Ethiopia. Proceeding of the first Ethiopian crop protection symposium, IAR, Addis Ababa, Ethiopia. pp. 79–108.

Esmail S. M., and Szabo, L. J. 2018. In: BGRI 2018 Technical Workshop. https://www.globalrust.org/content/wheat-stem-rust-pathogen-pgt-identification-and-characterizationegypt-using-single).

FAOSTAT, 2018. Agriculture Organization of the United Nations. Statistical Database. http://faostat.fao.org

Fetch Jr TG, and Dunsmore KM. 2004. Physiologic specialization of Puccinia graminis on wheat, barley, and oats in Canada in 2001. Canadian Journal of Plant Pathology, 26(2): 148–155.

Fetch, T. G., Park, R. F., Pretorius, Z. A., & Depauw, R. M. (2021). Stem rust: its history in Kenya and research to combat a global wheat threat. Canadian Journal of Plant Pathology, 43(sup2), S275–S297.

Gao, L., Babiker, E. M., Nava, I. C., Nirmala, J., Bedo, Z., Lang, L., … & Bariana, H. (2019). Temperature-sensitive wheat stem rust resistance gene Sr15 is effective against Puccinia graminis f. sp. tritici race TTKSK. Plant Pathology, 68(1), 143–151.

Gebrekirstos T, Woldeab G, and Salvaraj T. 2016. Distribution, Physiologic Races and Reaction of Wheat Cultivars to Virulent Races of Leaf Rust in Southeastern Zone of Tigray, Ethiopia. Regional Wheat Research or Development.

Getaneh Woldeab. 1996. Studies on fungal diseases of wheat at the Plant Protection Research Center, 1974-1994. In: Eshetu B., Abdurahman A. and Aynekulu Y. (eds.). Proceedings of the third annual conference of the Crop Protection Society of Ethiopia. Addis Ababa, Ethiopia. CPSE. Pp. 171–177.

Gutu, K., Tesfaye, T., Bacha, N., Negash, T., Kassa, D., Yirga, F., Debela, M., Bedada, G., Dessale, T., Gemechu, A. and Bayisa, T., 2022. Physiological Races of Puccinia graminis f. sp. tritici in Ethiopia in 2019/2020. American Journal of Agriculture and Forestry, 10(2), pp.72–76. Ethiop. J. Agric. Sci. 30(1) 87-97 (2020)

Hailu E, Woldaeb G, Danbali W, Alemu W, Abebe T (2015) Distribution of Stem Rust (Puccinia graminis f. sp. tritici) Races in Ethiopia. Adv Crop Sci Tech 3: 173. doi:10.4172/2329-8863.1000173

Hailu E, Woldaeb G, Denbel W, Alemu W, Abebe T, and Mekonnen A. 2015. Distribution of Stem Rust (Puccinia graminis f. sp. tritici) Races in Ethiopia. Plant Science 3(2): 15–19.

Hei N, Tsegaab T, Woldeab G, Hailu E, Bekele Hundie, Daniel Kassa, Fikirte Yirga, Fufa Anbessa, Wubishet Alemu, Teklay Abebe, Miruts Legesse, Alemar Seid, and Tesfaye Gebrekirstos. 2018. Distribution and frequency of wheat stem rust races (Puccinia graminis f. sp. tritici) in Ethiopia. Journal of Agricultural and Crop Research. 6(5): 88–96

Hei NB, Tsegaab T, Getaneh W, Girma T, Obsa C, Seyoum A, Zerihun E, Nazari K, Kurtulus E, Kavaz H, and Ozseven I. 2020. First Report of Puccinia graminis f. sp. tritici Race TTKTT in Ethiopia. Plant Disease, 104(3): 982.

Jaleta M, Hodson DP, Abeyo B, Yirga C, Erenstein O. Smallholders coping mechanisms with wheat rust epidemics: Lessons from Ethiopia. PLoS ONE. 2019; e0219327 10.1371/journal.pone.0219327

Jin Y, Szabo LJ, Pretorius ZA, Singh RP, Ward R, and Fetch Jr, T. 2008. Detection of virulence to resistance gene Sr24 within race TTKS of Puccinia graminis f. sp. tritici. Plant Disease, 92(6): 923–926.

Jin, Y., et al. 2008. Plant Dis. 92:923–926.

Kolmer JA. 2005. Tracking wheat rust on a continental scale. Current opinion in plant biology, 8(4): 441–449.

Lemma A, Woldeab G, Semahegn Y, and Dilnesaw Z. 2014. Survey and virulence distribution of wheat stem rust (Puccinia graminis f. sp. tritici) in the major wheat growing areas of central Ethiopia. Sci-Afric Journal of Scientific Issues, Research and Essays, 2(10): 474–478.

Li, F., Upadhyaya, N. M., Sperschneider, J., Matny, O., Nguyen-Phuc, H., Mago, R., … & Figueroa, M. (2019). Emergence of the Ug99 lineage of the wheat stem rust pathogen through somatic hybridisation. Nature communications, 10(1), 5068.

Luo, M., Xie, L., Chakraborty, S., Wang, A., Matny, O., Jugovich, M., et al. (2021). A five-transgene cassette confers broad-spectrum resistance to a fungal rust pathogen in wheat. Nat. Biotechnol. 39, 561–566. doi: 10.1038/s41587-020-00770-x

Mao, H., Jiang, C., Tang, C., Nie, X., Du, L., Liu, Y., … & Wang, X. (2023). Wheat adaptation to environmental stresses under climate change: Molecular basis and genetic improvement. Molecular Plant, 16(10), 1564–1589.

Mapuranga, J., Zhang, N., Zhang, L., Liu, W., Chang, J., & Yang, W. (2022). Harnessing genetic resistance to rusts in wheat and integrated rust management methods to develop more durable resistant cultivars. Frontiers in Plant Science, 13, 951095.

Meyer, M., Bacha, N., Tesfaye, T., Alemayehu, Y., Abera, E., Hundie, B., Woldeab, G., Girma, B., Gemechu, A., Negash, T. and Mideksa, T., 2021. Wheat rust epidemics damage Ethiopian wheat production: A decade of field disease surveillance reveals national-scale trends in past outbreaks. PloS one, 16(2), p.e0245697.

Mohamed, M. A., El Afandi, G. S., & El-Mahdy, M. E. S. (2022). Impact of climate change on rainfall variability in the Blue Nile basin. Alexandria Engineering Journal, 61(4), 3265–3275.

Mohammed, E. A., Zhi, X., & Abdela, K. A. (2025). Extreme weather patterns in Ethiopia: analyzing extreme temperature and precipitation variability. Atmosphere, 16(2), 133.

Nazari K, Kurtulus E, Kavaz H, Ozturk OM, Egerci Y, Cer C, Jarrahi T, Gasmi C, Kadiroglu A. First Report of Races TKTTP and TKKTP of Puccinia graminis f. sp. tritici with Virulence to Wheat Stem Rust Resistance Gene Sr24 in Turkey and Tunisia. Plant Dis. 2022 Feb;106(2):757. doi: 10.1094/PDIS-03-21-0450-PDN. Epub 2022 Jan 27. PMID: 34463527.

Olivera Firpo PD, Newcomb M, Flath K, Sommerfeldt-Impe N, Szabo LJ, Carter M, Luster DG, and Jin Y. 2017. Characterization of Puccinia graminis f. sp. tritici isolates derived from an unusual wheat stem rust outbreak in Germany in 2013. Plant pathology, 66(8): 1258–1266.

Olivera Firpo PD, Newcomb M, Szabo LJ, Rouse MN, Johnson JL, Gale SW, et al. Phenotypic and genotypic characterization of race TKTTF of Puccinia graminis f. sp. tritici that caused a wheat stem rust epidemic in southern Ethiopia in 2013/2014. Phytopathology. 2015; 105: 917–928. 10.1094/PHYTO-11-14-0302-FI [PubMed] [CrossRef] [Google Scholar]

Olivera P, Newcomb M, Szabo LJ, Rouse M, Johnson J, Gale S, Luster DG, Hodson D, Cox JA, Burgin L, Hort M, Gilligan CA, Patpour M, Juatesen AF, Hovmoller MS, Woldeab G, Hailu E, Hundie B, Tadese K, Pumphrey M, Sigh RP, and Jin Y. 2015. Phenotypic and Genotypic Characterization of Race TKTTF of *Puccinia graminis* f. sp. *tritici* that Caused a Wheat Stem Rust Epidemic in Southern Ethiopia in 2013 – 14. Phytopathology, 105(7): 917–928.

Olivera, P. D. et al. 2019. Phytopathology (10.1094/PHYTO-06-19-0186-R.

Olivera, P. D., Sikharulidze, Z., Dumbadze, R., Szabo, L. J., Newcomb, M., Natsarishvili, K., … & Jin, Y. (2019). Presence of a sexual population of Puccinia graminis f. sp. tritici in Georgia provides a hotspot for genotypic and phenotypic diversity. Phytopathology, 109(12), 2152–2160.

Patpour M, Afshari F, HasanBayat Z, Nazari K. 2014. Pathotype Identification of *Puccinia graminis* f.sp. *tritici*, the Causal Agent of Wheat Stem Rust under Greenhouse Conditions. Seed and Plant Improvement Institute-SPII, pp:26.

Patpour M, Hovmøller MS, Hansen JG, Justesen AF, Thach T, Algaba JR, Hodson D, and Randazzo B. 2018. Epidemics of yellow and stem rust in Southern Italy 2016-2017. In BGRI 2018. Retrieved on April, 2021 (https://www.globalrust.org).

Patpour M, Hovmøller MS, Rodriguez-Algaba J, Randazzo B, Villegas D, Shamanin VP, Berlin A, Flath K, Czembor P, Hanzalova A, Sliková S, Skolotneva ES, Jin Y, Szabo L, Meyer KJG, Valade R, Thach T, Hansen JG and Justesen AF (2022) Wheat Stem Rust Back in Europe: Diversity, Prevalence and Impact on Host Resistance. Front. Plant Sci. 13:882440. doi:10.3389/fpls.2022.882440

Patpour, M., Hovmøller, M.S., Justesen, A.F., Newcomb, M., Olivera, P., Jin, Y., Szabo, L.J., Hodson, D., Shahin, A.A., Wanyera, R. and Habarurema, I., 2016. The emergence of virulence to SrTmp in the Ug99 race group of wheat stem rust, Puccinia graminis f. sp. tritici, in Africa. Plant Dis., 100(522

Patpour, M., Hovmøller, M.S., Rodriguez-Algaba, J., Randazzo, B., Villegas, D., Shamanin, V.P., Berlin, A., Flath, K., Czembor, P., Hanzalova, A. and Sliková, S., 2022. Wheat stem rust back in Europe: Diversity, prevalence, and impact on host resistance. Frontiers in plant science, 13.

Patpour, M., Justesen, A.F., Tecle, A.W., Yazdani, M., Yasaie, M. and Hovmøller, M.S., 2020. First report of race TTRTF of wheat stem rust (*Puccinia graminis* f. sp. *tritici*) in Eritrea. Plant Disease, 104(3), p.973.

Pretorius ZA, Bender CM, Visser B, Terefe T. First Report of a Puccinia graminis f. sp. tritici Race Virulent to the Sr24 and Sr31 Wheat Stem Rust Resistance Genes in South Africa. Plant Dis. 2010 Jun;94(6):784. Doi: 10.1094/PDIS-94-6-0784C. PMID: 30754342.

Pretorius ZA, Bender CM, Visser B, Terefe T. First Report of a *Puccinia graminis* f. sp. *tritici* Race Virulent to the Sr24 and Sr31 Wheat Stem Rust Resistance Genes in South Africa. Plant Dis. 2010 Jun;94(6):784. Doi: 10.1094/PDIS-94-6-0784C. PMID: 30754342.

Rettie, F. M., Gayler, S., Weber, T. K., Tesfaye, K., & Streck, T. (2023). High-resolution CMIP6 climate projections for Ethiopia using the gridded statistical downscaling method. Scientific data, 10(1), 442.

Roelfs AP, and Groth JV, 1988. *Puccinia graminis* f. sp. *tritici*, black stem rust of Triticum spp. In Genetics of Plant Pathogenic Fungi, Advances in Plant Pathology, Vol. 6 (Sidhu, G.S., ed.). London: Academic Press, pp. 345–361.

Roelfs AP, RP Singh and EE Saari 1992. Rust Diseases of Wheat: Concept and Methods of Disease Management. Mexico, D.F.: CIMMYT, pp. 81.25

Saari E. E., and Prescott J. 1985. World distribution about economic losses. Mexico: CIMMYT.

Sanders R. Strategies to reduce the emerging wheat stripe rust disease. Synthesis of a dialog between policy makers and scientists from 31 countries at: the international Wheat Stripe Rust Symposium. Aleppo, Syria: International Center for Agricultural Research in the Dry Areas (ICARDA); 2011.

Silva, J. V., Reidsma, P., Baudron, F., Jaleta, M., Tesfaye, K., & van Ittersum, M. K. (2021). Wheat yield gaps across smallholder farming systems in Ethiopia. Agronomy for Sustainable Development, 41(1), 12.

Simane, B., Beyene, H., Deressa, W., Kumie, A., Berhane, K., & Samet, J. (2016). Review of climate change and health in Ethiopia: status and gap analysis. Ethiopian Journal of Health Development, 30(1), 28–41.

Singh RP, Hodson DP, Jin Y, Lagudah ES, Ayliffe MA, Bhavani S, Rouse MN, Pretorius ZA, Szabo LJ, Huerta-Espino J, Basnet BR, Lan C, Hovmøller MS. Emergence and Spread of New Races of Wheat Stem Rust Fungus: Continued Threat to Food Security and Prospects of Genetic Control. Phytopathology. 2015 Jul;105(7):872–84. doi: 10.1094/PHYTO-01-15-0030-FI. Epub 2015 Jun 29. PMID: 26120730.

Singh, B. K., Delgado-Baquerizo, M., Egidi, E., Guirado, E., Leach, J. E., Liu, H., & Trivedi, P. (2023). Climate change impacts on plant pathogens, food security and paths forward. Nature Reviews Microbiology, 21(10), 640–656.

Singh, R. P., et al. 2006. CAB Reviews: Perspectives in Agriculture, Veterinary Science, Nutrition and Natural Resources. 1. No. 054. CAB, Wallingford, U.K

Singh, R. P., Hodson, D. P., Jin, Y., Lagudah, E. S., Ayliffe, M. A., Bhavani, S., … & Hovmøller, M. S. (2015). Emergence and spread of new races of wheat stem rust fungus: continued threat to food security and prospects of genetic control. Phytopathology, 105(7), 872–884.

Sperschneider, J., Jones, A. W., Nasim, J., Xu, B., Jacques, S., Zhong, C., Upadhyaya, N. M., Mago, R., Hu, Y., Figueroa, M., Singh, K. B., Stone, E. A., Schwessinger, B., Wang, M. B., Taylor, J. M., & Dodds, P. N. (2021). The stem rust fungus Puccinia graminis f. sp. tritici induces centromeric small RNAs during late infection that are associated with genome-wide DNA methylation. BMC biology, 19(1), 203. 10.1186/s12915-021-01123-z

Stakman EC, Steward DM, and Loegering WQ. 1962. Identification of physiologic races of Puccinia graminis var. tritici. US Dep. Agric. Agric. Res. Serv. E-617.

Stubbs RW, Prescott JM, Sarrri EE, Dubin HJ. 1986. Cereal Disease Methodology Manual. CIMMYT, El Batan, Mexico, p.51

Teklay Abebe, Getaneh Woldeab, Woubit Dawit. 2012. Distribution and Physiologic Races of Wheat Stem Rust in Tigray, Ethiopia. Journal of Plant Pathology & Microbiology 3 (6): 142. doi:10.4172/2157-7471.1000142

Woldeab Getaneh, Hailu Endale, and Bacha Netsanet. 2017. Protocols for Race Analysis of Wheat Stem Rust (Puccinia graminis f. sp. tritici). Ethiopian Institute of Agricultural Research, Ambo Plant Protection Research Center, Ethiopia. Pp: 27.

Temam Hussien and DA Solomatin. 1984. Distribution of physiologic races of wheat stem rust in major growing regions of Ethiopia during 1982-83. Proceedings of the 9th Annual Meeting of Ethiopian Phytopathological Committee. Nazareth, Ethiopia. pp. 67–71.

Tesfaye, K., Tamene, L. D., Demissie, T. D., Seid, J., Haile, A., Mekonnen, K., & Solomon, D. (2021). A Framework for Bundling Climate-Smart Agriculture (CSA) and Climate Information Services (CIS) in Ethiopia.

Tofu, D. A., & Mengistu, M. (2023). Observed time series trend analysis of climate variability and smallholder adoption of new agricultural technologies in west Shewa, Ethiopia. Scientific African, 19, e01448.

Tsegaab, T., Alemayehu, C., Elfinesh, S., Hodson, D. and Szabo, L.J., 2020. First report of TTRTF race of wheat stem rust, Puccinia graminis F. sp. tritici, in Ethiopia. Plant Disease, 104(1), pp.293–293.

Tsushima, A., Lewis, C. M., Flath, K., Kildea, S., and Saunders, D. G. O. 2022. Wheat stem rust was recorded for the first time in decades in Ireland. Plant Pathol. 71, 890–900. doi: 10.1111/ppa.13532 (1)

Velásquez, A. C., Castroverde, C. D. M., & He, S. Y. (2018). Plant–pathogen warfare under changing climate conditions. Current biology, 28(10), R619–R634.

Wolday A, Fetch T, Hodson DP, Cao W, Briere S., 2011. First Report of Puccinia graminis f. sp. tritici Races with Virulence to Wheat Stem Rust Resistance Genes Sr31 and Sr24 in Eritrea.Plant Dis. 2011 Dec;95(12):1591. doi: 10.1094/PDIS-07-11-0582. PMID: 30732016

Wubaye, G. B., Gashaw, T., Worqlul, A. W., Dile, Y. T., Taye, M. T., Haileslassie, A.,… & Srinivasan, R. (2023). Trends in rainfall and temperature extremes in Ethiopia: station and agro-ecological zone levels of analysis. Atmosphere, 14(3), 483.

Wulff, B. B. H., and Krattinger, S. G. (2022). The long road to engineering durable disease resistance in wheat. Curr. Opin. Biotechnol. 73, 270–275. doi: 10.1016/j.copbio.2021.09.002

Yehizbalem Azmeraw, Belayneh Admassu, Bekele Abeyo, and Netsanet Bacha, 2020. Virulence Spectrum of Puccinia graminis f. sp. tritici in Northwest Ethiopia.DOI: 10.31038/AFS.2022433

Zadoks, J.C. and Bouwman, J.J. (1985) Epidemiology in Europe. In The Cereal Rusts, Vol. 2 (A.P. Roelfs and W.R. Bushnell, eds). Orlando, FL: Academic Press, pp. pp. 329–369.

Admassu, B., Lind, V., Friedt, W., and Ordon, F. (2009). Virulence analysis of Puccinia graminis f. sp. tritici populations in Ethiopia with special consideration of Ug99. Plant Pathology, 58(2), 362–369.

